# A novel approach to exploring the dark genome and its application to mapping of the vertebrate virus ‘fossil record’

**DOI:** 10.1101/2023.10.17.562709

**Authors:** Daniel Blanco-Melo, Matthew A. Campbell, Henan Zhu, Tristan P.W. Dennis, Sejal Modha, Spyros Lytras, Joseph Hughes, Anna Gatseva, Robert J. Gifford

## Abstract

**Background:** Genomic regions that remain poorly understood, often referred to as the “dark genome,” contain a variety of functionally relevant and biologically informative genome features. These include endogenous viral elements (EVEs) - virus-derived sequences that can dramatically impact host biology and serve as a virus “fossil record”. In this study, we introduce a database-integrated genome screening (DIGS) approach to investigating the dark genome *in silico*, focusing on EVEs found within vertebrate genomes.

**Results:** Using DIGS on 874 vertebrate species genomes, we uncovered approximately 1.1 million EVE sequences, with over 99% originating from endogenous retroviruses or transposable elements that contain EVE DNA. We show that the remaining 6038 sequences represent over a thousand distinct horizontal gene transfer events across ten virus families, including some that have not previously been reported as EVEs. We explore the genomic and phylogenetic characteristics of non-retroviral EVEs and determine their rates of acquisition during vertebrate evolution. Our study uncovers novel virus diversity, broadens knowledge of virus distribution among vertebrate hosts, and provides new insights into the ecology and evolution of vertebrate viruses.

**Conclusions:** We comprehensively catalogue and analyse EVEs within 874 vertebrate genomes, shedding light on the distribution, diversity and long-term evolution of viruses, and revealing their extensive impact on vertebrate genome evolution. Our results demonstrate the power of linking a relational database management system to a similarity search-based screening pipeline for *in silico* exploration of the dark genome.

## INTRODUCTION

The availability of whole genome sequence (WGS) data from a broad range of species provides unprecedented scope for comparative genomic investigations [1–3]. However, these investigations rely to a large extent on *annotation* - the process of identifying and labelling genome features - which usually lags far behind the generation of sequence data. Consequently, most whole genome sequences are comprised of DNA that is incompletely understood in terms of its evolutionary origins and functional significance. The portion of sequenced genome space that lacks annotations is sometimes referred to as the ‘dark genome’ [4], and contains a wide variety of yet-to-be-characterized genome features. Some of these may have functional roles, such as encoding proteins [5] or regulating gene expression [6]. Others, such as non-expressed pseudogenes, may not, but can nonetheless provide valuable insights into genome biology and evolution.

Within the dark genome, endogenous viral elements (EVEs) constitute a particularly intriguing group of genome features. EVEs are virus-derived DNA sequences that become integrated into the germline genome of host species and are stably inherited as host alleles – a form of horizontal gene transfer [7–14]. While once considered genetic ’junk,’ it has become evident over recent years that EVEs can profoundly impact host biology and genome evolution, with many now known to have physiologically relevant roles [15–19]. In addition, EVE sequences (whether functional or not) provide a rare source of retrospective information about ancient viruses, akin to a viral ‘fossil record’ [7, 20–22].

Identifying genome features contained within the dark genome, such as EVEs, often relies on the use of sequence similarity searches, such as those implemented in the Basic Local Alignment Search Tool (BLAST) [23, 24], to search WGS databases. Because sequence similarity reflects homology (evolutionary relatedness), novel genome features can often be identified based on their resemblance to ones that have been described previously. One example of this approach is implemented in the PSI-BLAST [5] and HMMR [8] programs, in which iterated search strategies are used to progressively increase sensitivity so that novel homologs of previously characterised genes may be detected. A related approach is ‘systematic *in silico* genome screening’ which extends the basic concept of a similarity search in two ways: (i) inclusion of multiple query sequences and/or target databases; (ii) similarity-based classification of matching sequences (‘hits’) via comparison to reference sequence library (**Fig. 1a**). Hits may also be further investigated using additional comparative or experimental approaches (**Fig. 1b**, **Table 1**). Thus, screening can provide one component of a broader analytical pipeline.

**Figure 1.**
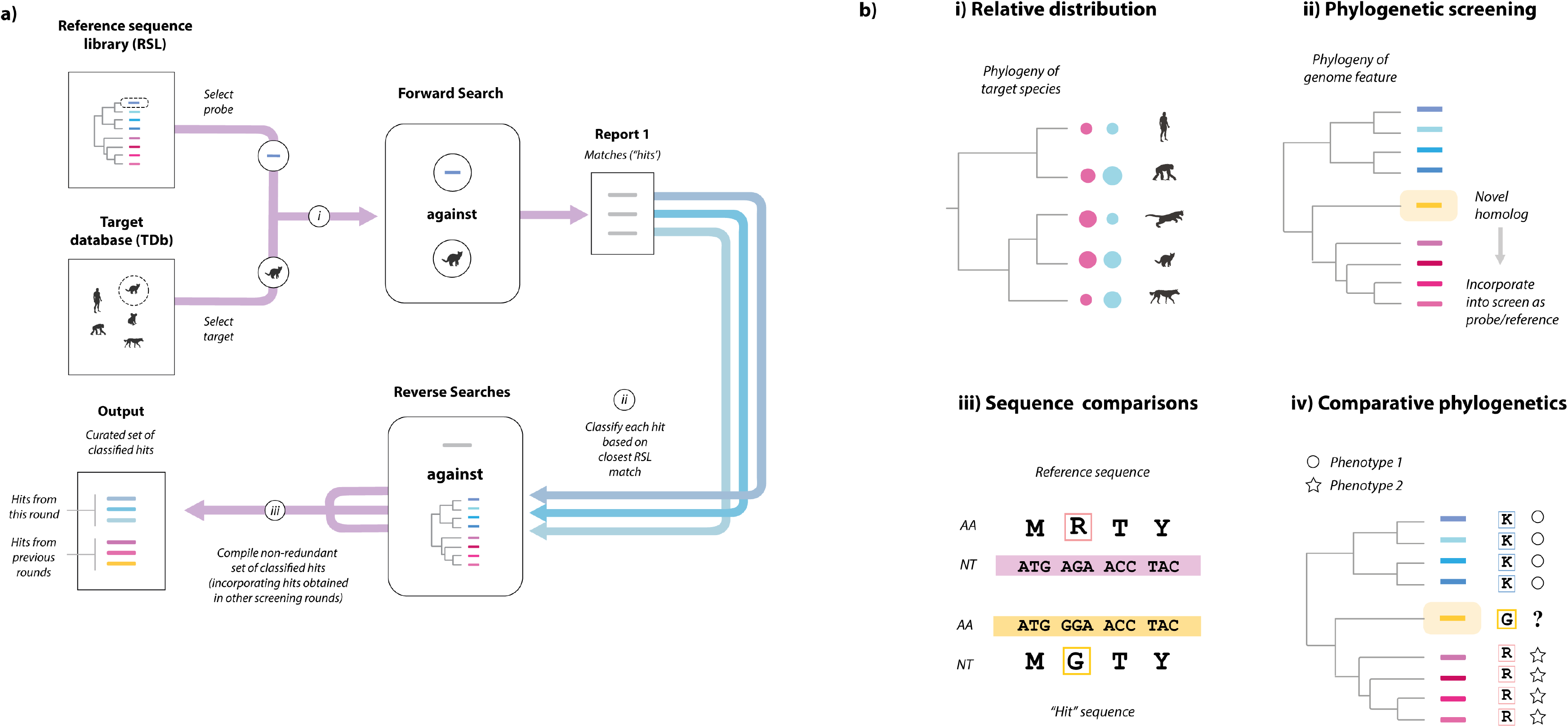
Exploring the dark genome using in silico screening. **(a) Overview of sequence similarity search-based screening.** Screening aims to identify and classify sequences similar to a set of query sequences within a target database (TDb) comprising whole genome sequence assemblies of multiple species. The schematic shows the steps that comprise a single round of screening, as follows: (i) A BLAST search is performed using a probe sequence selected from a curated ‘reference sequence library’ (RSL) and a ‘target’ file is selected from the TDb; (ii) Matching sequences (referred to as ‘hits’) identified in this screen are classified via similarity search-based comparison to the RSL; (iii) A non-redundant set of classified hits is compiled, incorporating hits from previous rounds of screening. **(b) Comparative analysis of screen output**. Sequences recovered via screening can be investigated using a wide range of *in silico*, comparative approaches, as follows: (i) analysis of feature distribution – e.g. annotating host phylogeny to show frequency of occurrence (coloured circles); (ii) phylogenetic screening, in which sequences obtained via similarity search-based screening are investigated in phylogenetic reconstructions (e.g. to identify novel lineages not present in the RSL, as shown here); (iii) Pairwise sequence comparisons – these can be used to identify specific differences in sequence obtained via screening, relative to reference sequences; (iv) Comparative phylogenetic analysis - the genetic properties of novel homologs can be inferred via comparative analysis (e.g. pairwise comparisons), while their phenotypic properties can potentially be investigated experimentally (e.g., via transcriptome sequencing).

**Table 1.**
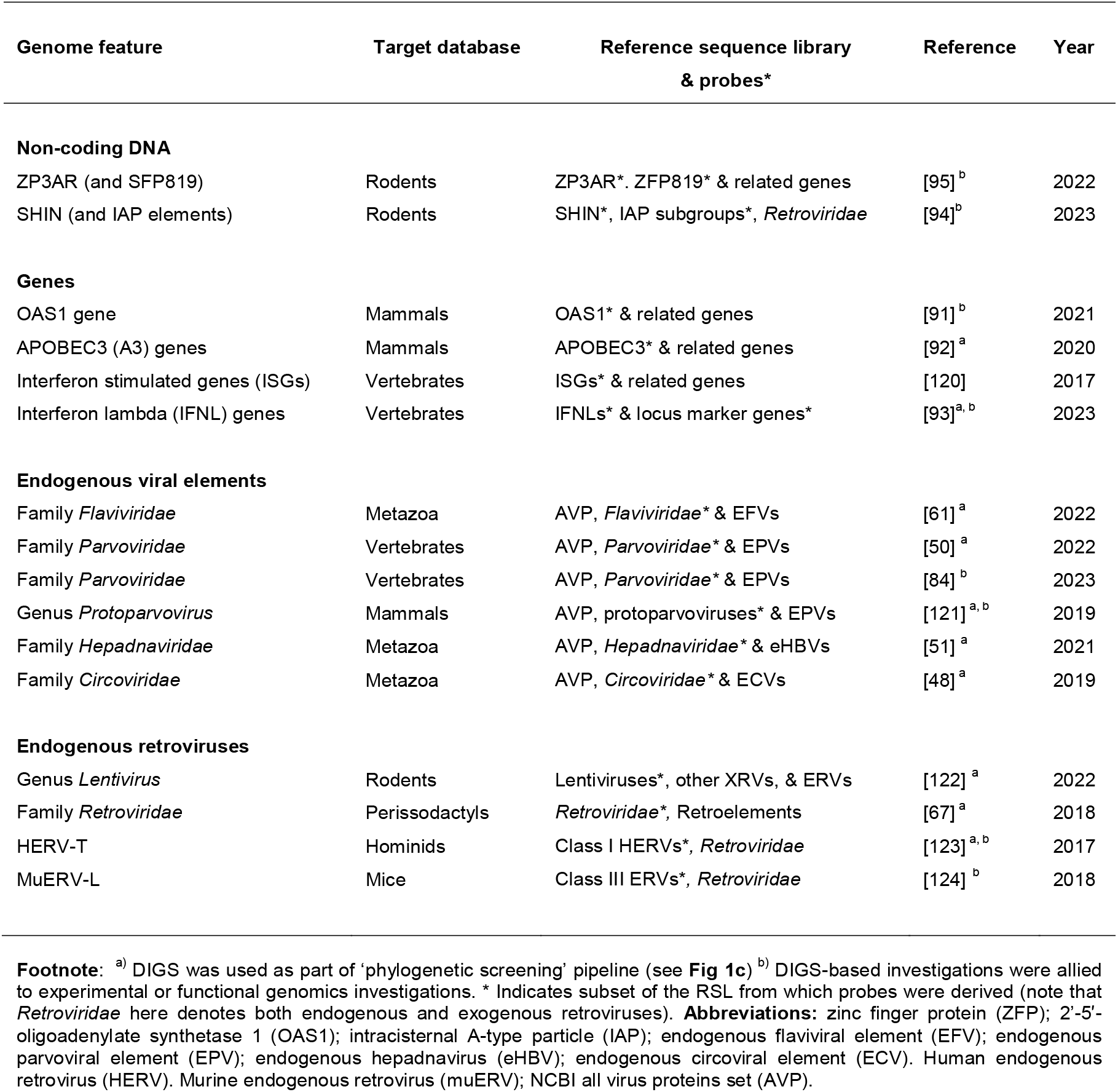
Examples of published studies utilising database-integrated screening.

While straightforward in principle*, in silico* genome screening is computationally expensive and can be difficult to implement efficiently. Moreover, large-scale screens can produce copious output data that are difficult to manage and interpret without an appropriate analytical framework. To address these issues, we developed a database-oriented approach to *in silico* screening, called *database-integrated genome screening* (DIGS). To demonstrate the use of this approach, we first created an open software framework for performing it, then used this framework to search published vertebrate genomes for EVE loci. Besides demonstrating that DIGS provides a powerful, flexible approach for exploring the dark genome, our analysis provides a comprehensive and detailed overview of EVE diversity in vertebrate genomes and reveals new information about the long-term evolutionary relationships between viruses and vertebrate hosts.

## RESULTS

### 1. A database-integrated approach to exploring the dark genome

We developed a robust, database-integrated approach to systematic *in silico* genome screening, referred to as database-integrated genome screening (DIGS). This approach integrates a similarity search-based screening pipeline with a relational database management system (RDBMS) to enable efficient exploration of the dark genome. The rationale for this integration is twofold: it not only provides a solid foundation for conducting large-scale, automated screens in an efficient and non-redundant manner but also allows for the structured querying of screening output using SQL, a powerful and well-established tool for database interrogation [25]. Additionally, an RDBMS offers advantages such as data recoverability, multi-user support, and networked data access.

The DIGS process comprises three key input data components:

**Target Database (TDb):** A collection of whole genome sequence assemblies (or other large sequence datasets such as transcriptomes) that will serve as the target for sequence similarity searches.

**Query Sequences (Probes):** A set of sequences to be used as input for similarity searches of the TDb.

**Reference Sequence Library (RSL):** The RSL represents the broad range of genetic diversity associated with the genome feature(s) under investigation. Its composition varies according to the analysis context (see **Table 1**). It should always include sequences representing diversity within the genome feature under investigation. It may also include genetic marker sequences and potentially cross-matching genome features. Probes are typically a subset of sequences contained in the RSL.

As illustrated in **Fig. 2**, the DIGS process involves systematic searching of a user-defined TDb with user-defined probes, merging fragmented hits, and classifying merged sequences through BLAST-based comparison to the RSL. The output - a set of non-redundant, defragmented ‘hits’ – is captured in a project-specific relational database. Importantly, this integration allows database queries to be employed in real time, with SQL queries referencing any information captured by the database schema. SQL-based querying of screening databases facilitates the identification of loci of interest, which can then be explored further using comparative approaches (see **Fig. 1b**).

**Figure 2.**
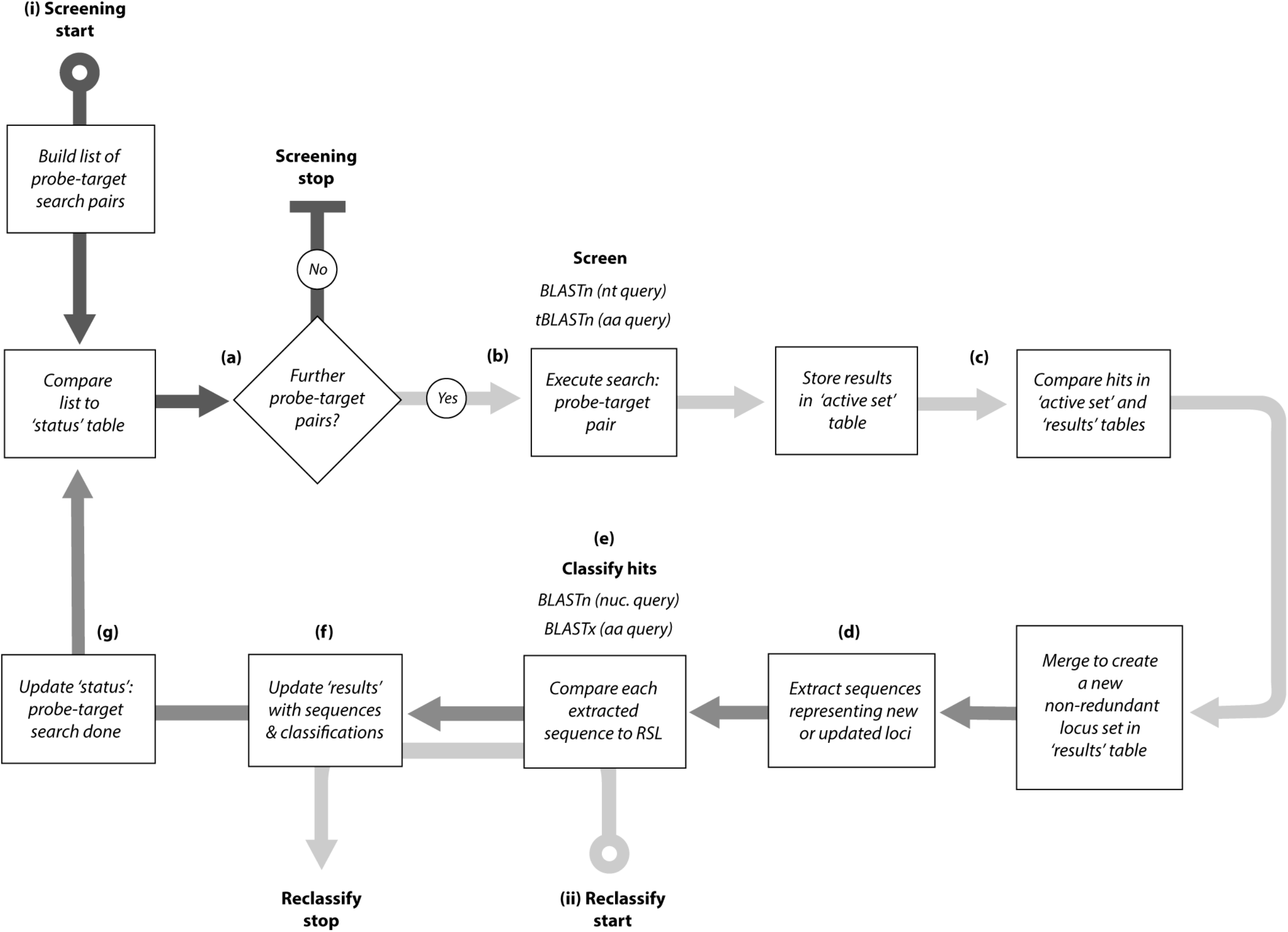
The database-integrated genome screening (DIGS) process as implemented in the DIGS tool. (i) Screening. **(a)** On initiation of screening a list of searches, composed of each query sequence versus each target database (TDb) file is composed based on the probe and TDb paths supplied to the DIGS program. Subsequently, screening proceeds systematically as follows: **(b)** the status table of the project-associated screening database is queried to determine which searches have yet to be performed. if there are no outstanding searches then screening ends, otherwise it proceeds to step **(b)** wherein the next outstanding search of the TDb is performed using the selected probe and the appropriate BLAST+ program. Results are recorded in the data processing table (‘active set’); **(c)** Results in the processing table are compared to those (if any) obtained previously to derive a non-redundant set of non-overlapping loci, and an updated set of non-redundant hits is created, with each hit being represented by a single results table row. To create this non- redundant set, hits that overlap, or occur within a given range of one another, are merged to create a single entry**; (d)** Nucleotide sequences associated with results table rows are extracted from TDb files and stored in the results table; **(e)** extracted sequences are classified via BLAST-based comparison to the RSL using the appropriate BLAST program; **(f)** The header-encoded details of the best-matching sequence (species name, gene name) are recorded in the results table. **(f)** The status table is updated to create a record of the search having been performed, and the next round of screening is initiated. **(ii)** Reclassification: Hits in the results table can be reclassified following an update to the reference sequence library.

It is important to note that screening is usually an iterative discovery process, wherein initial results inform the development of subsequent screens. For instance, novel diversity detected by an initial screen can subsequently be incorporated into the RSL, and hits within the screening database can be reclassified using the updated library (**Fig. 2**). Additionally, probe sets used in initial searches can be expanded to incorporate sequences identified during screening, broadening the range of sequences detected in subsequent screens [26]. However, care must be taken when using this approach, since it can potentially produce misleading results, or generate excessive hits (e.g. if highly repetitive sequences are contained within the new probes). Importantly, database integration allows results to be observed and interrogated in real time - as they are being generated. This means that configuration issues (e.g. badly composed RSL, inappropriate choice of probes) can be detected early on – potentially saving a significant amount of time and effort. Furthermore, it facilitates the implementation of agile, heuristic screening strategies, in which approaches are adjusted in line with results.

### 2. An open software framework for implementing DIGS

We constructed a software framework for implementing DIGS, called ‘the DIGS tool’. The DIGS tool is implemented using the PERL scripting language. It uses the BLAST+ program suite [24] to perform similarity searches, and the MySQL RDBMS (to capture their output). Accessible through a text-based console interface, it simplifies the complex process of large- scale genome screening, and provides a versatile basis for implementing screens.

To initiate screening using the DIGS tool, researchers provide a project-specific command file (**Fig. S1**) that serves as the blueprint for the screening process. This command file specifies parameters, including the user-defined name of the screening database, and file paths to the TDb, RSL, and probe sequences. When a screen is initiated a project-specific database is created with the schema shown in **Fig. S2**. This core schema can be extended to include any relevant “side data” – e.g., taxonomic information related to the species and sequences included in the screen - increasing the power of SQL queries to reveal informative patterns (**Fig. S2**, **Fig. S3**).

Systematic screening proceeds automatically until all searches have been completed. If the process is interrupted at any point, or if novel probe/target sequences are incorporated into the project, screening will proceed in a non-redundant way on restarting. Thus, screening projects can readily be expanded to incorporate new TDb files (e.g. recently published WGS assemblies), or novel probe/reference sequences, as they become available. The DIGS tool console allows reclassification of sequences held in the results table (e.g. following an RSL update). To increase efficiency, this process can be tailored to specific subsets of database sequences by supplying SQL constraints via the DIGS tool console.

BLAST algorithms emphasise local similarity and consequently tend to fragment contiguous matches into several separate hits if similarity across some internal regions of the match is low. The DIGS tool allows screening pipelines to be configured with respect to how overlapping/adjacent hits are handled, so that the screening process can be tailored to the specific needs of diverse projects. The DIGS tool also provides a ‘consolidation’ function that concatenates, rather than merges, adjacent hits and stores concatenated results, along with information about their structure, in a new screening database table.

For program validation, we mined mammalian genomes for sequences disclosing similarity to the antiviral restriction factor tetherin [27, 28]. Tetherin provides a useful test case as it is a relatively unique gene and its evolution, distribution and diversity have previously been examined [27, 28]. Results were compared with those provided by two alternative, widely used genome mining pipelines: OrthoDB [29] and Ensembl [30] and found to overlap by >99% (**Fig. S4**).

The DIGS tool provides functionality for exporting FASTA-formatted sequences and managing screening database tables (e.g., add/drop tables, import table data). Further information regarding program installation and usage is provided online, in a repository associated website [31]. In the sections below we illustrate the application of the DIGS tool to cataloguing of EVEs in vertebrate genomes, focussing on both high and low copy number elements.

### 3. Use of DIGS to catalogue RT-encoding endogenous retroviruses

Unlike other vertebrate viruses, retroviruses (family *Retroviridae*) integrate their genome into the nuclear genome of infected cells as an obligate part of their life cycle. As a result, retroviruses gain more opportunities to become a permanent part of the host germline. Furthermore, the initial integrated form of a retrovirus genome, called a provirus, is typically replication competent and increases in germline copy number can thus occur through reinfection of germ line cells [32]. Accordingly, ‘endogenous retroviruses’ (ERVs) are by far the most common type of EVE found in vertebrate genomes [7, 33].

Retrovirus genomes contain a *pol* coding domain that encodes a reverse transcriptase (RT) gene. The RT gene can be used to reconstruct phylogenetic relationships across the entire *Retroviridae* and hence provides the lynchpin for unravelling the evolutionary history and origins of ERV loci [34, 35]. We therefore implemented a screening procedure to detect RT- encoding ERV loci, based on an RSL comprised of previously classified exogenous retrovirus and ERV RT sequences (**see Methods**). Screening involved more than 1.5 million discrete tBLASTn searches and resulted in the identification of 1,073,137 ERV RT hits. This set was filtered based on higher BLAST bitscore cut-off to obtain a high confidence set of 500,701 loci (**Table 2**).

**Table 2.** ERV RT loci identified via *in silico* screening.

|  |  | Retrovirus clade |  |  |  |  |  |
| --- | --- | --- | --- | --- | --- | --- | --- |
| Vertebrate class | # WGS | Clade I |  | Clade II |  | Clade III |  |
|  |  | Total # | Average # | Total # | Average # | Total # | Average # |
| Agnatha | 3 | 32 | 10.67 | 1* | 0.33 | 300 | 100.00 |
| Chondrichthyes | 6 | 2018 | 336.33 | 0 | 0.00 | 2843 | 473.83 |
| Actinopterii | 173 | 8514 | 49.21 | 64* | 0.37 | 2177 | 12.58 |
| Actinistia | 1 | 0 | 0.00 | 0 | 0.00 | 97 | 97.00 |
| Amphibia | 34 | 17319 | 509.38 | 973 | 28.62 | 8019 | 235.85 |
| Reptiles | 92 | 13676 | 148.65 | 12120 | 131.74 | 20197 | 219.53 |
| Aves | 143 | 17951 | 125.53 | 20797 | 145.43 | 42014 | 293.80 |
| Mammalia | 452 | 13676 | 30.26 | 174549 | 386.17 | 143364 | 317.18 |

**Legend.** WGS=whole genome sequence assemblies screened. \* Hits likely due to contamination.

High confidence ERV RT hits were identified in all vertebrate classes. However, the frequency among classes was found to vary dramatically (**Fig. 3**). ERVs occur most frequently in mammals (class Mammalia) and amphibians (class Amphibia), and at relatively similar, intermediate frequencies in the genomes of reptiles (class Squamata) and birds (class Aves). By contrast, RT-encoding ERVs are relatively rare in the genomes of fish, including ray-finned fish (class Actinopterygii) and jawless fish (class Agnatha). Cartilaginous fish (class Chondrichthyes) represent a possible exception, although only a few genomes were available for this group (**Fig. 3**). These findings are broadly consistent with previous studies, conducted using a smaller amount of species genomes [33, 36–38].

**Figure 3.**
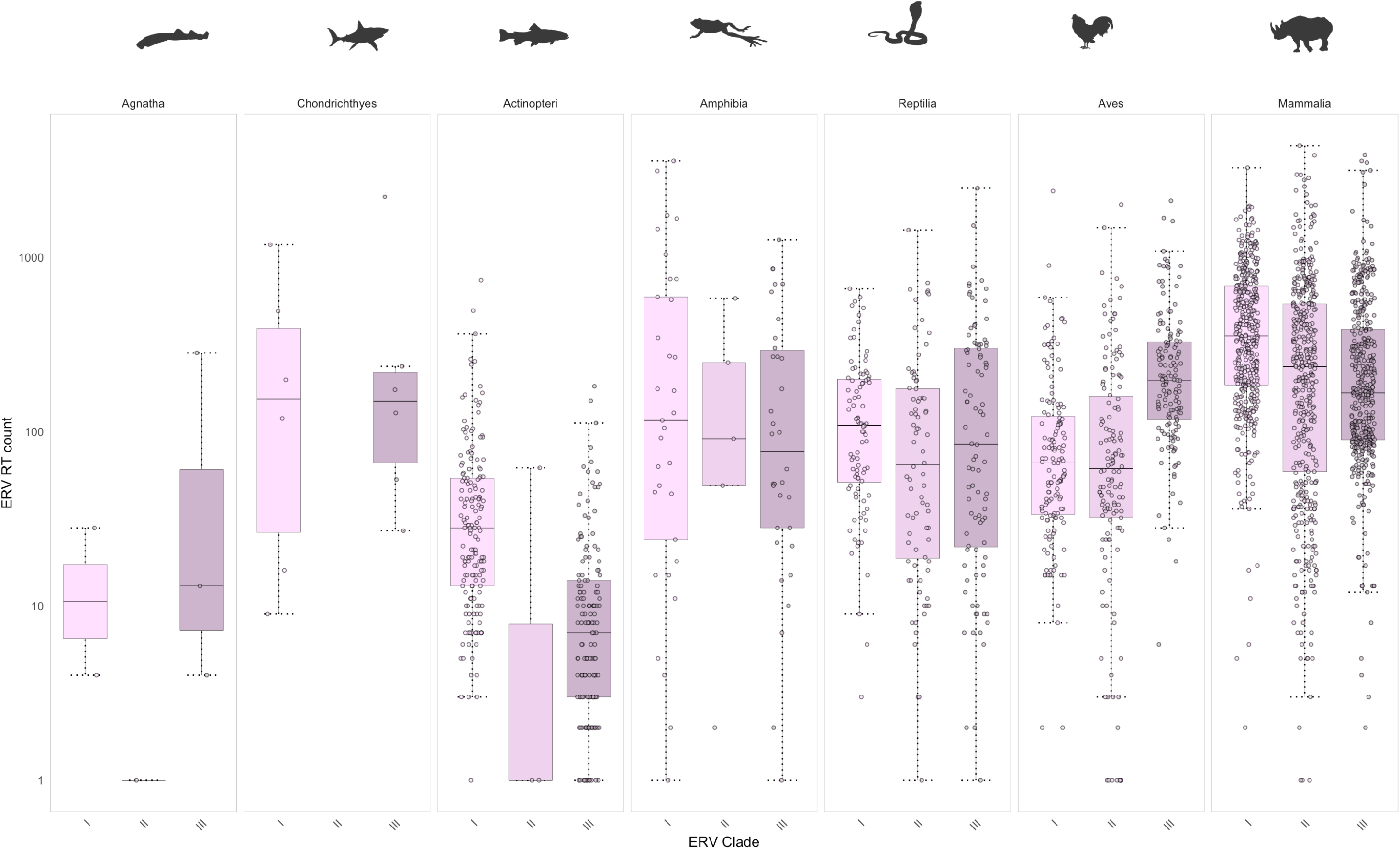
Counts of ERV RT loci identified by identified via database integrated genome screening of 874 vertebrate species. Box plots showing the distribution of endogenous retrovirus (ERV) reverse transcriptase (RT) counts in distinct vertebrate classes. Median and range of values are indicated. Circles indicate counts for individual species. Counts are shown against a log scale. Figure created in R using ggplot2 and geom_boxplot.

ERVs have been taxonomically grouped into three clades (I, II and III) based on their phylogenetic relatedness in the RT gene to the exogenous *Gammaretrovirus*, *Betaretrovirus* and *Spumavirus* genera respectively [1,2]. We incorporated into our RT screening database taxonomic information for (i) host species examined in our screen and (ii) RSL RT sequences. We then used SQL statements referencing these tables to summarise the frequency of clade I, II and III ERVs in distinct vertebrate classes. Whereas clade I and III ERVs are found in all vertebrate groups, clade II ERVs appear to have a more restricted distribution, occurring only at low frequency in amphibians, and being completely absent from agnathans and cartilaginous fish (**Table 2**). A few clade II ERVs were identified in ray-finned fish, but these were very closely related to mammalian ERVs and likely represent contamination of WGS builds with mammalian genomic DNA. While RT-encoding ERV copy number is relatively high in cartilaginous fish, RT diversity is relatively low, with the majority of ERV RT sequences belonging to clade III.

### 4. Use of DIGS to catalogue non-retroviral EVEs vertebrate genomes

To identify non-retroviral EVEs, we first obtained an RSL representing all known viruses [39]. From this library, a set of representative probes was selected. Probes include representative proteomes of all known vertebrate viruses except retroviruses. Screening entailed >1.5 million discrete tBLASTn searches, and initial results comprised 33,654 hits. However, many of these represented matches to host genes and TEs. We identified these spurious matches by interrogating screening results with a combination of SQL queries and *ad hoc* phylogenetic analysis. We also identified and excluded hits that appeared likely to derived from exogenous viruses (see **Table S1**). For example, SQL-generated summaries of our initial screen results revealed several WGS sequences disclosing unexpected similarity to plant virus genomes (**Fig. S3a**). Among these, matches to geminiviruses (family *Geminiviridae*) and potyviruses (family *Potyviridae*) lack evidence for germline integration and likely to represent diet-related contamination. Other unusual matches were contained within large contigs and thus could not be dismissed as contaminating DNA but were revealed to be spurious by genomic and phylogenetic analysis. For example, a sequence identified in the genome of the pig-nosed turtle (*Carettochelys insculpta*) disclosed similarity to caulimoviruses (family Caulimoviruses) - but was revealed by closer analysis to represent an unusual ERV (**Fig. S3a**, **Fig. S5**).

Next we removed matches to transposons partly comprised of virus-derived DNA such as the adintovirus-derived mavericks [40] and alloherpesvirus-derived teratorns [41] (**Fig. S3a**). Once TEs had been removed, results comprised 6038 putative non-retroviral EVE sequences, representing 10 virus families (**Table 3**, [42]). We did not identify any EVEs derived from vertebrate viruses with genomes comprised of double-stranded RNA (e.g., order Reovirales) or circular single-stranded RNA (e.g., genus *Deltavirus*). However, all other virus genome ‘classes’ were represented including reverse-transcribing DNA (DNArt) viruses, double-stranded DNA (DNAds) viruses, single-stranded DNA (DNAss) viruses, single-stranded negative sense RNA (RNAss-ve) viruses, and single-stranded positive sense RNA (RNAss+ve) viruses. Plotting the distribution of EVEs and exogenous viruses from distinct virus families and genera across vertebrate phyla, revealed that many virus groups have had a broader distribution across vertebrate hosts than recognised on the basis of exogenous isolates (**Fig. 4**).

**Figure 4.**
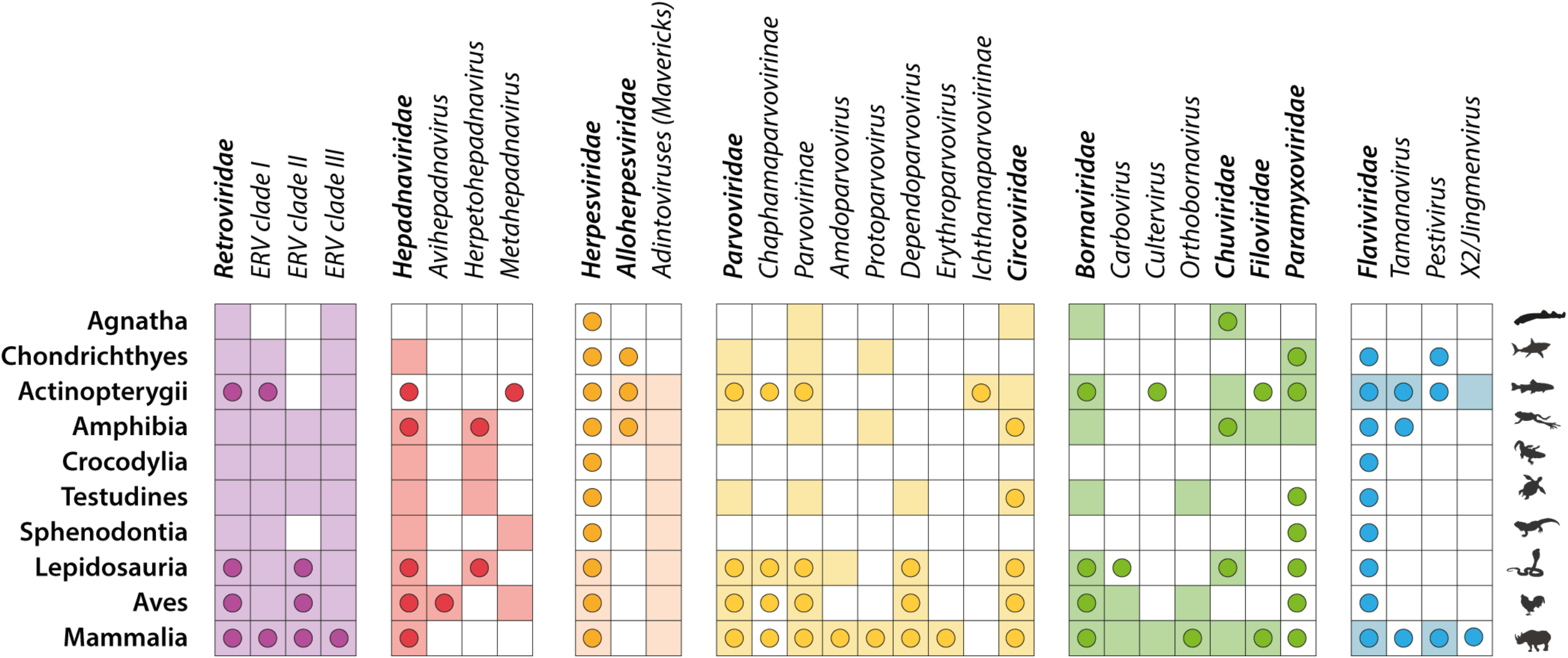
Distribution of virus families across vertebrate hosts. Circles indicate the presence of exogenous viruses. Shaded boxes indicate the presence of confirmed endogenous viral elements. Abbreviations: DNArt = reverse transcribing DNA viruses; DNAss = single-stranded DNA viruses; DNAds double-stranded DNA viruses; RNAds = double-stranded RNA viruses; RNAss-ve = single-stranded negative sense RNA viruses; RNAss+ve = RNAss-ve = single-stranded positive sense RNA viruses. Information on the distribution of exogenous viruses was obtained from the NCBI virus genomes resource [39], supplemented with information obtained from recently published papers [56, 113–118].

**Table 3.** Number of non-retroviral EVE sequence identified and estimated number of germline incorporation events in distinct vertebrate classes.

| Virus Group | # EVEs identified (Estimated # germline incorporation events) |  |  |  |  |  |  |  |  |  |  |  |  |  |  |  |
| --- | --- | --- | --- | --- | --- | --- | --- | --- | --- | --- | --- | --- | --- | --- | --- | --- |
|  | Total |  | Mammalia |  | Aves |  | Reptiles |  | Amphibia |  | Actinopterygii |  | Chondrichthyes |  | Agnatha |  |
| ssRNA-ve |  |  |  |  |  |  |  |  |  |  |  |  |  |  |  |  |
| Bornaviridae | 2566 | (383) | 2434 | (292) | 30 | (11) | 27 | (14) | 52 | (44) | 22 | (21) | - | - | 1 | (1) |
| Chuviridae | 182 | (164) | 24 | (24) | - | - | 23 | (23) | 9 | (9) | 119 | (108) | - | - | 7 | - |
| Filoviridae | 390 | (69) | 389 | (68) | - | - | - | - | 1 | (1) | - | - | - | - | - | - |
| Paramyxoviridae | 19 | (17) | - | - | - | - | - | - | 4 | (3) | 14 | (3) | 1 | (1) | - | - |
| ssRNA+ve |  |  |  |  |  |  |  |  |  |  |  |  |  |  |  |  |
| Flaviviridae | 8 | (11) | 1 | (1) | - | - | - | - | - | - | 7 | (10) | - | - | - | - |
| DNArt |  |  |  |  |  |  |  |  |  |  |  |  |  |  |  |  |
| Hepadnaviridae | 993 | (108) | - | - | 897 | (89) | 93 | (17) | 2 | (1) | - | - | 1 | (1) | - | - |
| DNAss |  |  |  |  |  |  |  |  |  |  |  |  |  |  |  |  |
| Circoviridae | 1198 | (131) | 918 | (29) | 32 | (15) | 91 | (19) | 82 | (24) | 68 | (38) | - | - | 7 | (6) |
| Parvoviridae | 689 | (238) | 534 | (199) | 34 | (10) | 34 | (13) | 12 | (7) | 12 | (6) | 3 | (3) | - | - |
| DNAds |  |  |  |  |  |  |  |  |  |  |  |  |  |  |  |  |
| Herpesviridae | 13 | (8) | 11 | (6) | 1 | (1) | 1 | (1) | - | - | - | - | - | - | - | - |
| Alloherpesviridae | 28 | (8) | - | - | - | - | - | - | 15 | (3) | 13 | (5) | - | - | - | - |
| Total | 6087 | (1137) | 4311 | (619) | 994 | (126) | 269 | (87) | 177 | (92) | 255 | (191) | 5 | (5) | 15 | (7) |

**Legend:** Germline incorporation here implies both integration into the germline and fixation.

We examined all EVE loci identified in our study to determine their coding potential. We identified numerous EVE loci encoding open reading frames (ORFs) >300 amino acids (aa) in length (**Fig. S6**). Among these, four encoded ORFs longer than 1000 aa. One of these – a 1718aa ORF encoded by an endogenous borna-like L-protein (EBLL) element in bats (EBLL-Cultervirus.29-EptFus) – has been reported previously [43]. However, we also identified an endogenous chuvirus-like L-protein (ECLL) element encoding an ∼1400 aa ORF in livebearers (subfamily Poeciliinae). This element encodes long ORFs in two distinct livebearer species (*P. formosa and P. latapina*), indicating it’s coding capacity has been conserved for >10 million years [44]. We also detected herpesvirus and alloherpesvirus EVEs encoding ORFs >1000 aa, but as discussed below, the integration status of these sequences remains unclear.

### 5. Diversity of non-retroviral EVEs in vertebrate genomes

#### 5.1 EVEs derived from viruses with double-stranded DNA genomes

We detected DNA derived from herpesviruses (family *Herpesviridae*) in mammalian and reptilian genomes (**Fig. 4**, **Table 3**, [42]). DNA sequences derived from betaherpesviruses (subfamily *Betaherpesvirinae*) and gammaherpesviruses (subfamily *Gammaherpesvirinae*) have previously been reported in WGS assemblies of the tarsier (*Carlito syrichta*) and aye- aye (*Daubentonia madagascensis*), respectively [45]. In addition to these sequences, we detected gammaherpesvirus DNA in WGS data of red squirrels (*Sciurus vulgaris*) and the Amazon river dolphin (*Inia geoffrensis*), while betaherpesvirus DNA was detected in the stoat (*Mustela ermina*) WGS assembly, and DNA derived from an alphaherpesvirus (subfamily *Alphaherpesvirinae*) in the Oriximina lizard (*Tretioscincus oriximinensis*) WGS (**Fig. S7a-b**). Germline integration of human betaherpesviruses has been demonstrated [46, 47], and the presence of a betaherpesvirus-derived EVE in the tarsier genome EVE has been established [45]. However, herpesviruses can also establish latent infections, and might be considered likely to occur as contaminants of DNA samples used to generate whole genome sequence assemblies. Due to the limitations of the WGS assemblies in which they were identified, it was not possible to confirm that the novel herpesvirus DNA sequences detected here represent EVEs rather than DNA derived from contaminating exogenous viruses.

DNA derived from alloherpesviruses (family *Alloherpesviridae*) was detected in fish and amphibians. In ray-finned fish, most of these sequences belonged to the ’teratorn’ lineage of transposable elements, which have arisen via fusion of alloherpesvirus genomes and piggyBac transposons, and have been intragenomically amplified in the genomes of teleost fish (Infraclass Teleostei) [41]. Additional alloherpesvirus-related elements were identified in three amphibian species and five ray-finned fish species [42]. One of these elements, identified in the Asiatic toad (*Bufo gargarizans*) occurred within a contig that was significantly larger than a herpesvirus genome, demonstrating that it represents an EVE rather than an exogenous virus. Phylogenetic analysis revealed that alloherpesvirus-like sequences identified in amphibian genomes clustered robustly with amphibian alloherpesviruses, while those identified in fish genomes clustered with fish alloherpesviruses (**Fig. S7c**).

#### 5.2 EVEs derived from viruses with single-stranded DNA genomes

EVEs derived from parvoviruses (family *Parvoviridae*) and circoviruses (family *Circoviridae*) are widespread in vertebrate genomes, being found in the majority of vertebrate classes (**Fig. 4**). Both endogenous circoviral elements (ECVs) and endogenous parvoviral elements (EPVs) are only absent in major vertebrate groups represented by a relatively small number of sequenced species genomes (i.e. between 1 and 6). No ECVs or EPVs were identified in the tuatara (order Rhynchocephalia) or in crocodiles (order Crocodilia). EPVs were not identified in agnathans, while ECVs were not identified in cartilaginous fish.

We identified a total of 1192 ECVs, most of which derived from an element in carnivore (Class Mammalia: order Carnivora) genomes that is embedded within a non-LTR retrotransposon and has undergone intragenomic amplification (**Fig. S8**). While many of the

ECVs identified in our screen have been reported in previous publications [7, 26, 48–50], we also identified novel loci in mammals, reptiles, amphibians, and ray-finned fish [42]. Phylogenetic analysis (see **Fig. S7d**) revealed that a novel ECV locus in turtles was found to group with avian circoviruses, while amphibian ECV elements grouped with fish circoviruses, though bootstrap support for this relationship was lacking. A circovirus-like sequence detected in the WGS data of Allen’s wood mouse (*Hylomyscus alleni*) grouped robustly with the exogenous ‘rodent circovirus’, but integration of this sequence into the *H. alleni* genome could not be confirmed.

We identified 627 EPVs, representing two distinct subfamilies within the *Parvoviridae* and five distinct genera (see **Fig. 4**). The majority of these loci have been reported in a previous study of vertebrate genomes [50] or were related to these loci. However, we also identified novel EPVs in reptiles, amphibians and mammals (**Table 3**, [42]). In reptiles the novel elements derived from genus *Dependoparvovirus* while the amphibian elements were more closely related to viruses in genus *Protoparvovirus*. Interestingly, these EPVs clustered basally within a clade of protoparvovirus-related viruses in phylogenetic reconstructions (**Fig. S7e**), consistent with previous analyses indicating that this genus may have broadly co-diverged with vertebrate phyla [50].

#### 5.3 EVEs derived from reverse-transcribing DNA viruses

EVEs derived from hepadnaviruses (family *Hepadnaviridae*), which are reverse-transcribing DNA viruses, were identified in reptiles, birds and amphibians (**Table 3**, [42]). Most of these EVEs, which are commonly referred to as ‘endogenous hepatitis B viruses’ (eHBVs), have been reported previously [51, 52]. However, we identified novel elements in the plateau fence lizard (*Sceloporus tristichus*), and also in vertebrate classes where they have not been reported previously. These include one eHBV identified in a cartilaginous fish, the Australian ghostshark (*Callorhinchus milii*), and another identified in an amphibian, the common coquí (*Eleutherodactylus coqui*).

Phylogenetic analysis (see **Fig. S7f**) revealed that novel eHBV elements identified in lizards (suborder Lacertilia) group robustly with the exogenous skink hepadnavirus (SkHBV), while the amphibian element groups with robustly within a clade comprised of the exogenous spiny lizard hepadnavirus (SlHBV), Tibetan frog hepadnavirus (TfHBV) and eHBV elements identified in crocodile genomes. The eHBV identified in sharks was relatively short and not amenable to phylogenetic analysis, but nonetheless provides the first evidence that hepadnaviruses infect this host group.

#### 5.4 EVEs derived from viruses with single-stranded, negative sense RNA genomes

Screening revealed that vertebrate genomes contain numerous EVEs derived from mononegaviruses (order *Mononegavirales*), which are characterised by non-segmented ssRNA-ve genomes. These EVEs derive from four mononegavirus families: bornaviruses (family *Bornaviridae*), filoviruses (family *Filoviridae*), paramxyoviruses (family *Paramyxoviridae*) and chuviruses (family *Chuviridae*) (**Fig. 4**, **Table 3**, [42]). We did not detect any EVEs derived from other mononegavirus families that infect vertebrates (*Pneumoviridae*, *Rhabdoviridae*, *Nyamiviridae*, *Sunviridae*), nor any EVEs derived from virus families with segmented, negative sense RNA genomes (e.g., *Peribunyaviridae*, *Orthomyxoviridae*).

The majority of mononegavirus EVEs identified in our screen were derived from bornaviruses and filoviruses and have been described in previous reports [7, 48, 50, 51, 53]. However, we also identified novel EVEs derived from these groups, as well as previously unreported EVEs derived from paramyxoviruses and chuviruses (**Table 3**). Germline integration of DNA derived from mononegaviruses can occur if, in an infected germline cell, viral mRNA sequences are reverse transcribed and integrated into the nuclear genome by cellular retroelements [54]. EVE loci generated in this way preserve the sequences of individual genes of ancient mononegaviruses, but not entire viral genomes. Among mononegavirus-derived EVEs, regardless of which family, EVEs derived from the nucleoprotein (NP) and large polymerase (L) genes predominate, but other genes are also represented, including the glycoprotein (GP) genes of filoviruses, bornaviruses, and chuviruses, the VP30 and VP35 genes of filoviruses, and the hemagglutinin-neuraminidase (HA-NM) gene of paramyxoviruses.

Paramyxovirus-like EVEs were identified in ray-finned fish, amphibians and sharks (**Fig. 4**, **Table 3**, [42]). Many of these EVEs were highly divergent and/or degenerated and consequently their evolutionary relationships to contemporary paramyxoviruses were poorly resolved in phylogenetic analysis. However, an L polymerase-derived sequence identified in the pobblebonk frog (*Limnodynastes dumerilii*) genome was found to group robustly with sunshine virus, a contemporary paramyxovirus of Australian pythons [55] in phylogenetic trees (**Fig. S7g**).

Chuvirus-like sequences were identified in agnathans, ray-finned fish, reptiles, and mammals (**Fig. 4**, **Table 3**, [42]). The majority of the mammalian elements were identified in marsupials, but we also identified a single chuvirus-like EVE in the genome of a laurasiatherian mammal – the bottlenose dolphin (*Tursiops truncatus*). Phylogenetic trees reconstructed using alignments of NP-derived chuvirus EVEs and NP genes of contemporary chuviruses revealed evidence for the existence of distinct clades specific to particular vertebrate classes (**Fig. S7h**). These included a clade including both a snake EVE and an exogenous chuviruses of snakes, and two clades comprised of EVEs and viruses of teleost fish. In addition, these phylogenies revealed a robustly supported relationship between chuvirus EVEs in the Tibetan frog (*Nanorana parkeri*) and zebrafish (*Danio rerio*) genomes. Taken together, these results provide evidence for the existence of numerous diverse lineages of chuviruses in vertebrates, adding to recent evidence for the presence of exogenous chuviruses in marsupials [56].

Filovirus-derived EVEs were mainly identified in mammals (**Fig. 4**, **Table 3**, [42]). However, we also identified one filovirus-derived EVE in an amphibian – the mimic poison frog (*Ranitomeya imitator*) - providing the first evidence that filoviruses infect this vertebrate group (**Table 1**). We identified novel, ancient filovirus EVEs in anteaters (family Myrmecophagidae) and spiny mice (genus *Acomys*).

Strikingly, the inclusion of Tapajos virus (TAPV), a snake filovirus, in phylogenetic reconstructions revealed evidence for the existence of two highly distinct filovirus lineages in mammals (**Fig. 5**). These two lineages, which are robustly separated from one another by TAPV, are evident in phylogenies constructed for both the NP and VP35 genes. One lineage (here labelled ‘Mammal-1’) is comprised of EVEs and all contemporary mammalian filoviruses, whereas the other (‘Mammal-2’) is comprised exclusively of EVEs. Notably, within the Mammal-1 group, EVEs identified in mammals indigenous to Southern Hemisphere continents (e.g. marsupials, xenarthrans) group basally, whereas EVEs and viruses isolated from ‘Old World’-associated placental mammals occupy a more derived position.

**Figure 5.**
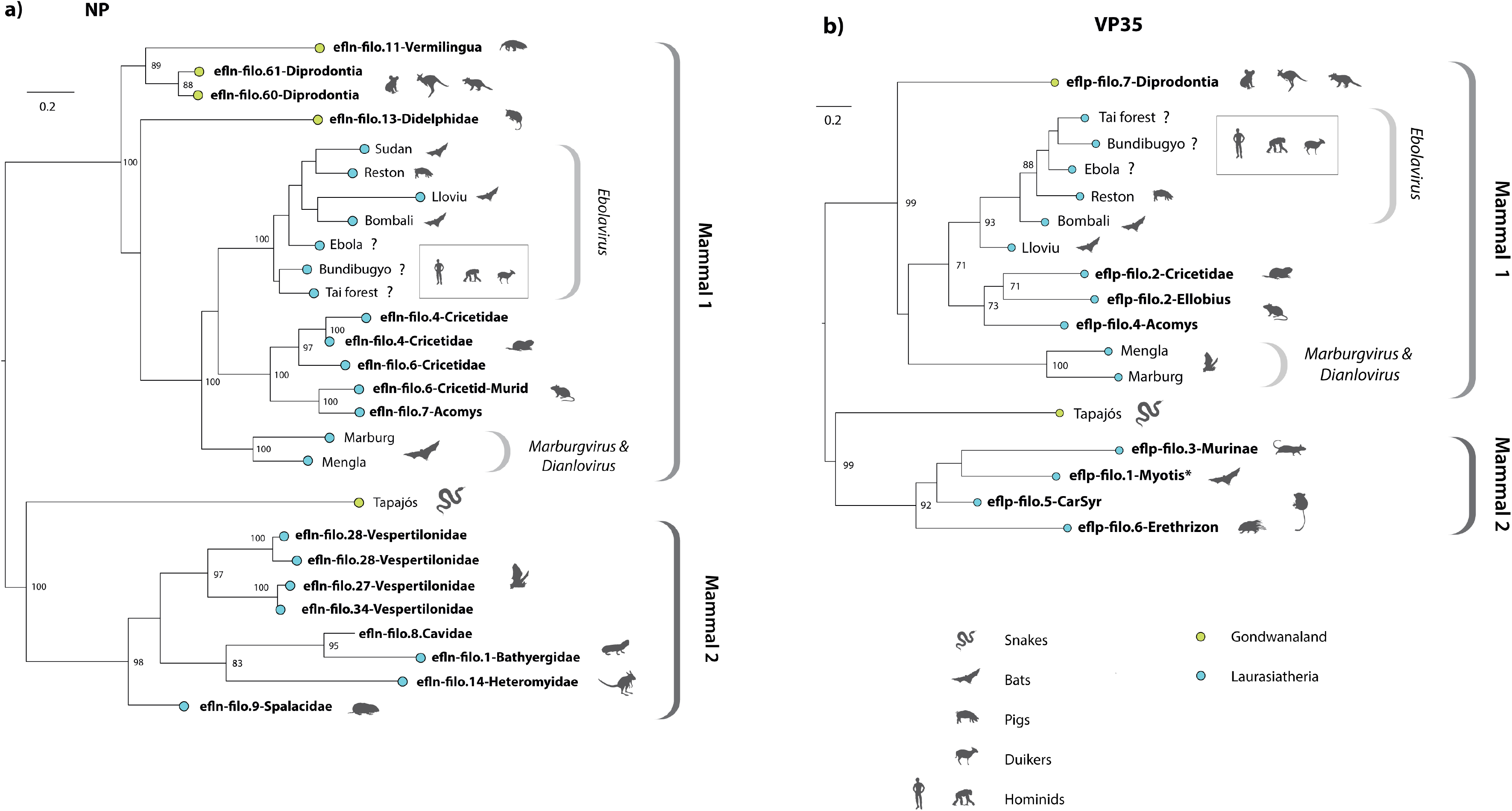
Evolutionary relationships of filoviruses and filovirus-derived EVEs. Bootstrapped maximum likelihood phylogenies showing the evolutionary relationships between filoviruses and filovirus EVEs in the nucleoprotein (NP) and viral protein 35 (VP35) genes. Phylogenies were constructed using maximum likelihood as implemented in RAxML, and codon-aligned nucleotides for each gene. Numbers adjacent internal nodes indicate bootstrap support (1000 bootstrap replicates). The scale bar indicates evolutionary distance in substitutions per site. Virus taxon names are shown in regular font, EVE names are shown bold. EVE names follow standardised nomenclature (see Methods). Brackets to the right of each tree indicate virus genera (italics) and major lineages (bold). Silhouettes indicate host groups following the key. For Ebola virus, Bundibugyo virus, and Tai Forest virus, the main reservoir hosts are unknown. The inset box adjacent these taxa shows host species in which one or more of these viruses has been isolated [73, 119], following the key.

The ‘Mammal-2’ clade contains filovirus EVEs from rodents, primates and bats. Because EVEs belonging to this clade were obtained from several distinct lineages, and show conservation across these groups, we can be reasonably confident they represent a *bona fide* lineage within the *Filoviridae*, rather than just a set of highly degraded filo-like EVEs that group together due to long branch attraction [57]. One member of this group (eflp-filo.1- Myotis) encodes an intact VP35 protein, the properties of which have been experimentally investigated in recent studies [58, 59]. Interestingly, we found that spiny mice also harbour a filovirus EVE encoding an intact VP35 protein (eflp-filo.3-Acomys), however, this insertion belongs to the ‘Mammal 1’ clade and is relatively closely related to the VP35 proteins found in contemporary mammalian filoviruses (**Fig. 5b**).

Bornavirus-like EVEs were identified in all vertebrate classes except Chondrichthyes (**Fig. 4**, **Table 3**, [42]). The majority have been reported previously or are orthologs of previously reported EVEs. However, we identified novel bornavirus-like EVEs in the genomes of ray-finned fish and amphibians. The amphibian EVEs grouped robustly with culterviruses in phylogenetic reconstructions (**Fig. S7i-j**).

#### 5.5 EVEs derived from viruses with single-stranded, positive sense RNA genomes

EVEs derived from positive sense RNA viruses are rare in vertebrate genomes (**Fig. 4**, **Table 3**, [42]). The only examples we identified were a small number of sequences derived from flavivirids (family *Flaviviridae*). These include an EVE derived from the *Pestivirus* genus of flavivirids, the reference genome of the Indochinese shrew (*Crocidura indochinensis*), as reported previously [60], and EVEs identified in ray-finned fish, also reported previously [61]. In fish genomes, flavivirid EVEs derive from the proposed ‘Tamanavirus’ genus, and a lineage labelled ‘X2’ that groups as a sister taxon to the proposed ‘Jingmenvirus’ genus. However, jingmenviruses are actually segmented, RNAss-ve viruses whose genomes include flavivirid-derived segments [62]. Since is possible that the X2 lineage shares a common RNAss-ve ancestor with jingmenviruses, EVEs belonging to this lineage may in fact be derived from viruses with ssRNA-ve genomes.

### 6. Frequency of germline incorporation events across distinct vertebrate phyla

We used the DIGS framework to dissect the history of horizontal gene transfer events involving germline incorporation of DNA derived from non-retroviral viruses. We excluded EVEs derived from DNAds viruses from this analysis because most of these are mavericks/polintons or teratorn elements that have undergone intragenomic amplification. For these groups, the large number of insertions, and the fact that amplified lineages appear to have been independently established on multiple occasions, meant that such an analysis would be beyond the scope of this study. Furthermore, for most of the few dsDNA-derived EVEs that did not belong to these groups, it was not possible to determine if they represented germline-integrated elements, exogenous viruses, or integrations occurring in somatic cells.

To examine the rate of germline incorporation in the remaining groups of non-retroviral EVEs representing DNAss, DNArt, RNAss-ve and RNAss+ve viruses, we compiled an expanded RSL containing a single reference sequence for each putative (or previously confirmed) ortholog. By classifying our hits against this expanded RSL, we could discriminate novel EVE loci (paralogs) from orthologs of previously described EVE loci. Where novel paralogs were identified, we incorporated these into our RSL and then reclassified related sequences in our screening database against this updated library. By investigating loci in this way, and iteratively reclassifying database sequences, we progressively resolved the various non- retroviral EVEs identified in our screen into sets of putatively orthologous insertions. Via this analysis we estimated that the non-retroviral EVEs identified in our study (excluding those derived from dsDNA viruses) represent ∼1137 distinct germline incorporation events (**Table 3**). Using orthology information we calculated minimum age estimates for all non-retroviral EVEs identified in two or more species [42]. We applied standardised nomenclature to EVE loci (see **Methods**), capturing information about EVE orthology, taxonomy, and host distribution [42].

Next, we estimated the rate of germline incorporation for each endogenized virus family, in all vertebrate classes represented by at least ten species (**Fig. 6**). Rates were found vary dramatically across each of the vertebrate groups examined. Overall, rates were highest in mammals, and lowest in reptiles. Fish and amphibians disclosed similar rates with ssDNA and ssRNA-ve viruses being incorporated at similar, intermediate rates. Birds were generally similar to reptiles but show a higher rate of ssDNA virus incorporations and a markedly elevated rate of hepadnavirus incorporation. Rates of parvovirus, filovirus, and bornavirus infiltration were very high in mammals compared to other vertebrate classes, with bornaviruses being incorporated at a particularly high rate (>0.03 per million years of species evolution). A relatively high rate of incorporation of RNAss+ve viruses was observed in ray- finned fish, but since the elements in question are closely related to jingmenviruses, as described above, they may in fact reflect incorporation of DNA derived from an RNAss-ve virus group [62].

**Figure 6.**
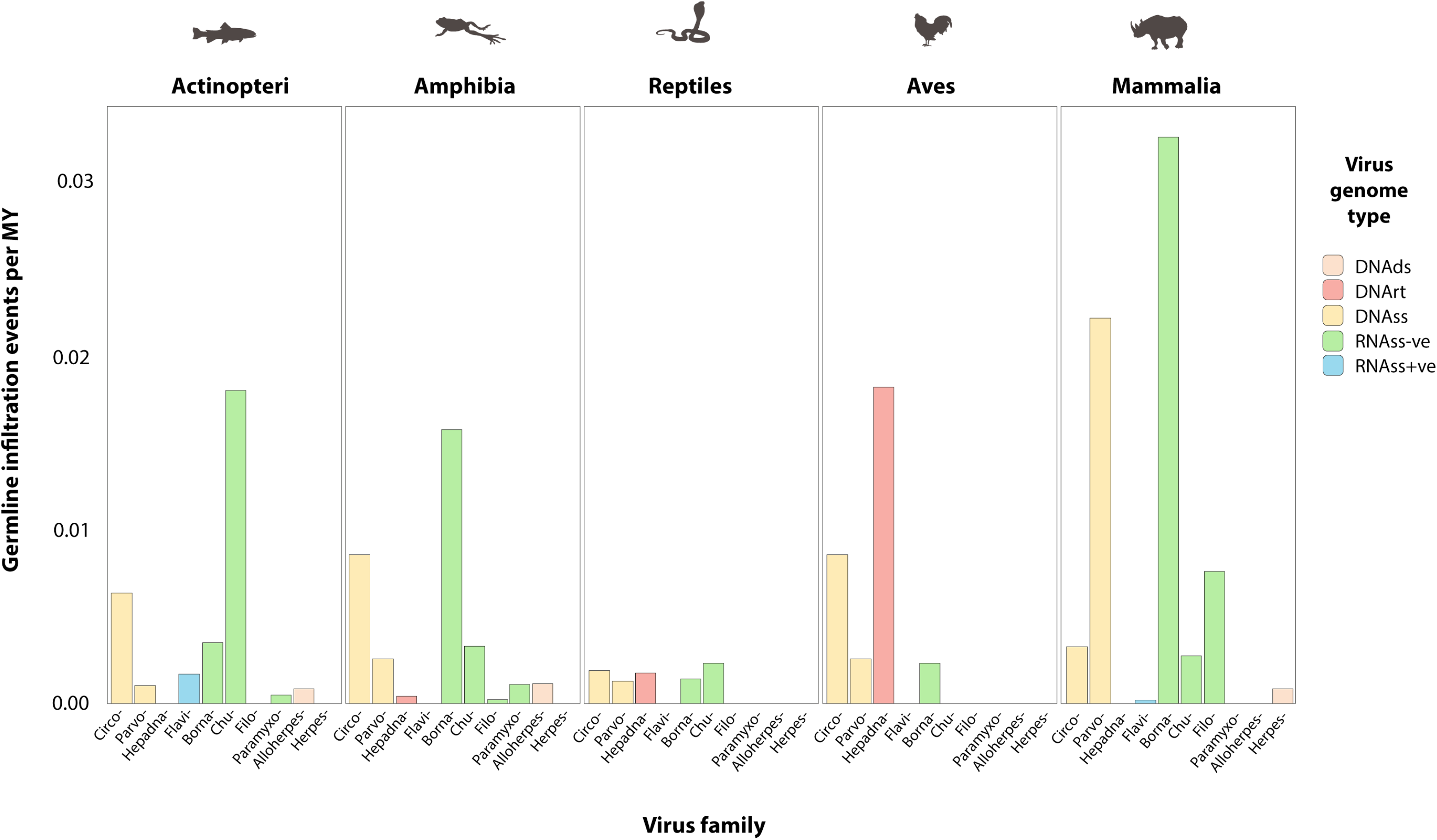
Comparison of germline infiltration rates in five vertebrate classes. Infiltration rates represent the rate of incorporation and fixation per million years (MY) of species branch length sampled. Rates are shown for each non-retroviral family represented by vertebrate EVEs. Colours indicate reverse transcribing DNA (DNArt) viruses, single stranded DNA (DNAss) viruses, single stranded negative sense RNA (RNAss-ve) virueses and single stranded positive sense RNA (RNAss-ve) viruses, following the key.

In addition to estimating the frequency of germline incorporation of non-retroviral viruses, we used our screening data to reconstruct a time-calibrated overview of virus integration throughout vertebrate evolutionary history (**Fig. 7, Table S2**). Among putatively orthologous groups of EVEs for which we were able to estimate minimum dates of integration, the majority were found to have been incorporated in the Cenozoic Era (1-66 Mya). So far, the oldest integration event identified involves a metahepadnavirus (genus *Metahepadnavirus*)- derived EVE that appears to be orthologous in tuataras and birds, indicating it was incorporated into the saurian germline >280-300 Mya (see [51]). Other ancient EVEs include circovirus and herpetohepadnavirus (genus *Herpetohepadnavirus*)-derived EVEs in turtles (order Testudines) (see [52]), a circovirus-derived EVE in frogs (order Anura), and bornavirus integrations in placental mammals (see [53]). Besides revealing the landscape of non-retroviral EVE integration throughout vertebrate history, plotting EVE distribution in this way clearly reveals the main differences in EVE distribution across host groups (**Fig. 7**).

**Figure 7.**
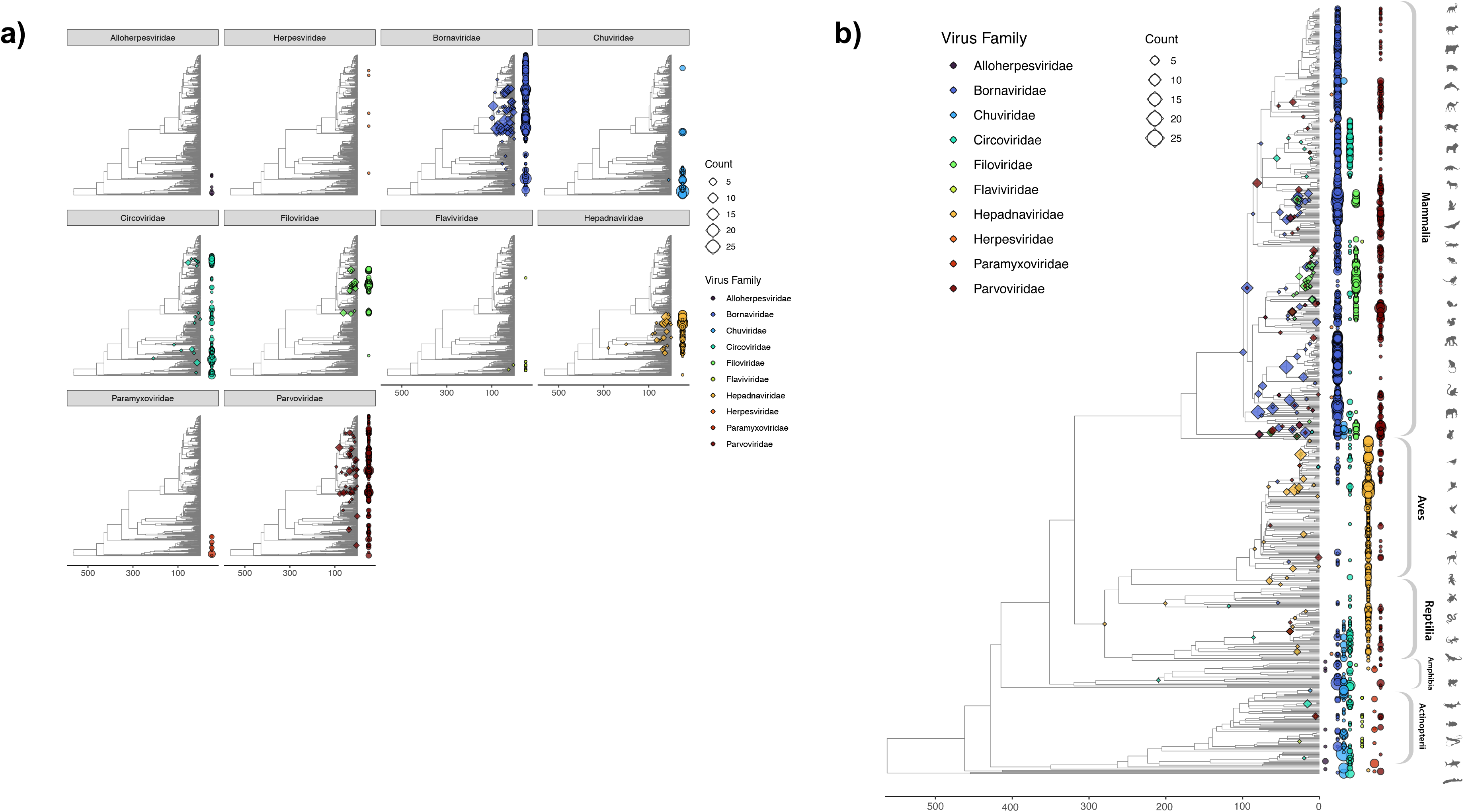
Overview of germline incorporation in vertebrates. A time-calibrated phylogeny of vertebrate species examined in this study, obtained via TimeTree [109]. Minimum ages of endogenization events are indicated by diamonds on internal nodes for EVE loci present as orthologs in multiple species. The presence of EVE sequences in each species genome is indicated by circles at phylogeny tips. Circles and diamonds nodes are scaled by the number of sequences detected and color-coded by virus family as indicated in legend. For circles, scaling indicates the total number of EVE sequences detected within each species genome, including both unique and shared endogenization events. In panel **(a)** the distribution of 10 families of viruses is shown across vertebrates separately. In panel **(b)** all virus families are shown on the same tree.

## DISCUSSION

Sequencing of genomes is advancing rapidly but deciphering the complex layers of information they contain is a challenging, long-term endeavour [58, 59]. Genomes are not only inherently complex, they also exhibit remarkable dynamism, with phenomena such as recombination, transposition and horizontal gene transfer contributing to the creation of genomic ‘churn’ that makes feature distribution difficult to map [60]. These issues, combined with rapid data accumulation, coverage limitations, and assembly errors – make generation of complete and accurate annotations difficult [62, 63]. Consequently, labour intensive manual genome annotation remains important [58, 61], and most published whole genome sequences are comprised of genomic ‘dark matter’.

An exciting aspect of these circumstances is that they provide immense scope to make interesting biological discoveries using low cost, *in silico* approaches. While experimental studies are generally required to characterise genome features at a functional level, approaches based solely on comparative sequence analysis (see **Fig 1b**) can often reveal useful insights into their biology and evolution [1, 63]. Furthermore, comparative investigations *in silico* can often be productively combined with functional genomics or experimental approaches (**Fig 1b**, **Table 1**).

Systematic *in silico* genome screening is computational approach that facilitates investigation of the dark genome (**Fig. 1**). However, it can be challenging to implement efficiently. Automated pipelines are generally required to implement large-scale screens [64], and these can produce copious output data that are difficult to manage and interpret without an appropriate analytical framework. In this report, we introduce DIGS – a robust analytical platform for conducting large-scale *in silico* screens - and describe an open software framework (the DIGS tool) for implementing it.

EVEs constitute one interesting and informative group of genome features that can be found within the dark genome [22]. They are poorly annotated for several reasons. Firstly, they arise sporadically via horizontal gene transfer, and consequently their distribution is unpredictable [7, 22]. Additionally, uncharacterised EVE loci may be hard to recognise due to their being highly degraded or fragmented, or because their exogenous virus counterparts are either unknown or extinct [65, 66]. Finally, there are numerous potential sources of confounding or artefactual results that can arise during EVE screening, including host genes that exhibit similarity to virus genes, and contamination of WGS assemblies with DNA derived from exogenous viruses.

To illustrate how DIGS facilitates identification and characterisation of features hidden within the dark genome, we use the DIGS tool to perform a broad-based investigation of EVE diversity in vertebrates. We first focussed on high-copy number EVEs - which in vertebrate genomes mainly comprise ERVs. We screened 874 vertebrate genomes for RT-encoding ERVs and identified 700,000 high confidence matches. This screen revealed marked differences in ERV RT copy number between vertebrate classes. An in-depth investigation of ERV diversity in vertebrates – for example, examining their composition in finer detail, or incorporating insertions that lack RT sequences, was considered beyond the scope of this study. However, the RT dataset generated here provides a robust foundation for further ERV studies that are underpinned by phylogenetic analysis. For example, we have previously used RT data in combination with other *in silico* approaches for in-depth, phylogenetical characterisation of ERVs within discrete mammalian subgroups (e.g. see [67]).

ERVs constitute a unique type of EVE, in that they can remain replication-competent following integration and may increase their germline copy number through continued virus replication. However, the germline copy number of any EVE can potentially increase through interactions with TEs - this has been described for ERVs [68–70], as well as for EVEs derived from dsDNA viruses [40, 41, 71]. In addition, data obtained here and in our previous investigations show that EVEs derived from hepadnaviruses have been amplified in cormorants [51], while circovirus-derived sequences have been amplified in carnivore genomes [48], apparently in association with LINE1 activity [42]. These findings underline the impact of fusion between EVEs and vertebrate transposons on vertebrate genome evolution. This phenomenon has occurred on multiple independent occasions and involved a diverse range of vertebrate viruses. Interestingly, we found that circovirus EVEs in carnivore genomes are associated with a retroelement lineage (LINE1) that has also inserted into gammaherpesvirus and Chikungunya virus genomes (**Fig. S8**). These findings suggest that retroelement-mediated transposition can establish a complex network of horizontal gene transfer events linking host and virus genomes.

DIGS is not only well-suited to exploring the distribution and diversity of high copy number genome features such as ERVs and TEs, it can also be used in ‘beach combing’ searches that aim to identify rare and unusual genome features. These kinds of screens typically require a rigorous filtering process to distinguish genuine from spurious matches, and as shown here, this is facilitated by database integration. DIGS enabled the efficient identification of EVEs derived from non-retroviral viruses (which are relatively rare and diverse) and provided a powerful framework for filtering spurious results (**Fig. S3**).

Via DIGS, we established a broad overview of non-retroviral EVE diversity in vertebrate genomes (**Fig. 4**, **Fig. 6**), shedding new light on virus distribution and diversity in vertebrates. Notably, our findings extend the known host range of important virus families. For example, we identify a filovirus-derived EVE in a frog (order Anura), providing the first evidence for the existence of amphibian filoviruses. In addition, we provide the first evidence for the presence (at least historically) of hepadnaviruses in sharks, and chuviruses in mammals (**Fig. 4**). In addition, we reveal novel virus diversity. For example, we identify novel lineages of parvoviruses and circoviruses in amphibians (**Fig S7d-e**), as well as a novel circovirus lineage in turtles (**Fig S7d**) and a novel hepadnavirus lineage in frogs (**Fig S7f**). We also identify novel paramyxovirus, chuvirus and bornavirus lineages in fish and amphibians (**Fig. S7g-j**).

Mammalian filoviruses include some of the most lethal viruses in the world [72], and while the natural reservoirs of some are known, they remain unclear for the highly pathogenic ebolavirus (EBOV) and its closest relatives (**Fig. 5**). EBOV is assumed to have a zoonotic origin, but it has rarely been possible to formally link outbreaks to a given animal reservoir, limiting understanding of its emergence. So far, efforts to identify the true reservoirs of ebolaviruses have tended to focus on bats [73]. However, the widespread presence of filovirus EVEs in rodents [42], including some groups that have not been examined as potential EBOV reservoirs, such as spiny mice, suggests that the potential of this group to serve as a reservoir should not be overlooked.

Previous studies have noted that filovirus EVE sequences in the genomes of cricetid rodents (family Cricetidae) robustly split the *Ebolavirus* and *Cuevavirus* genera from the *Marburgvirus* and *Dianlovirus* genera, demonstrating that these groups diverged >20 million years ago (Mya) [74], rather than within the past 10,000 years as suggested by molecular clock-based analysis of contemporary filovirus genomes [75]. Here, we found that TAPV, an exogenous virus of snakes, robustly separates two clades of mammalian filoviruses in phylogenetic reconstructions. Since transmission of filoviruses between reptiles and mammals is likely quite rare, and both lineages contain ancient EVEs (**Fig. 5, Table S2**), these findings support the long-term existence of two highly distinct filovirus lineages (‘mammal 1’ and ‘mammal 2’) in mammals. Notably, basal taxa within the ‘mammal 1’ lineage – which also includes all known contemporary filoviruses of mammals – disclose associations with Southern Hemisphere continents (Australia, South America) that were largely isolated throughout extensive periods of the Cenozoic Era. These data suggest that filoviruses were present in ancestral mammals inhabiting Gondwanaland (an ancient supercontinent comprised of South America, Africa, India, and Australia) and diversified into at least two major lineages as mammalian populations became compartmentalised in distinct continental regions during the early to mid-Cenozoic. An interesting question is whether the ‘mammal 2’ group represents filoviruses that evolved in Northern hemisphere- associated, boreoeutherian mammals (magnorder Boreoeutheria), while ‘mammal 1’ represents filoviruses that initially evolved in Southern hemisphere-associated marsupials (infraclass Marsupialia) and xenarthrans (magnorder Xenarthra) before disseminating throughout the globe (possibly in association with volant mammals – i.e., bats).

While several previous studies have described EVE diversity in vertebrates [33, 36, 76], our investigation is significantly larger in scale and breadth. Furthermore, for non-retroviral viruses, we introduced a higher level of order to EVE data, making use of the DIGS framework to discriminate orthologous versus paralogous EVE loci, and to identify intra- genomically amplified EVE lineages. This allowed us to establish a panoramic view of germline incorporation by non-retroviral viruses during vertebrate evolution (**Fig. 7**). Furthermore, discriminating orthologous and paralogous EVEs enabled us to infer the rates of germline infiltration by non-retroviral virus families with greater accuracy than in previous studies (**Fig. 6**, **Fig. 7**). Notably, we did not find strong evidence for a reduced rate of germline infiltration in avian genomes, as suggested by a previous study [77]. Incorporation of DNArt viruses is higher in birds than in any other vertebrate class (**Fig. 6**), while acquisition of EVEs derived from ssDNA-ve viruses does appear to be limited in this group, they closely resemble reptiles in this respect. Furthermore, avian hosts appear similar to reptiles with regard to ERV RT copy number (**Fig. 3**).

The absence, or near absence, of many virus groups from our catalogue of vertebrate EVEs is noteworthy. For example, vertebrates are infected with many ssRNA+ve viruses, but EVEs derived from these viruses are extremely rare, while EVEs derived from viruses with circular RNA genomes, or double-stranded RNA genomes, were not detected at all (**Table 3**). All other virus genome types were represented by EVEs in the vertebrate germline, but their distribution is patchy and limited to a relatively small number of virus families (**Fig. 4**, **Fig 7**). For example, among ssRNA-ve viruses, only mononegaviruses were found to be present – we found no evidence for germline integration of segmented ssRNA-ve viruses such as orthomyxoviruses and bunyaviruses. The scarcity of EVEs derived from these virus groups suggests that aspects of their biology strictly limit their capacity to for germline incorporation. These likely include cell tropism (whether germline cells are typically infected) and the site of cellular replication (with viruses replicating in the nucleus more likely to be incorporated). Additionally, vertebrate germline cells may present strong intrinsic barriers to the replication of certain virus groups.

The catalogue of EVE loci generated here provides a foundation for further investigations in virology, genomics, and human health. From the virology perspective, EVEs provide information about the long-term evolutionary history viruses, which greatly influences how we understand their biology. As well as enabling future studies of vertebrate ‘paleoviruses’, the EVE catalogue can inform efforts to identify and characterise new viruses (both by providing ecological and evolutionary insights [49], and by helping identify ’false positive’ hits arising from genomic DNA). From the genomics side, EVEs are of interest due to their important roles in physiology and genome evolution [78]. These include roles antiviral immunity [11, 79, 80], and a diverse range of other physiological processes [18, 58, 59, 81–83]. Notably, we identified numerous non-retroviral EVEs encoding ORFs longer than 300aa (**Fig. S6**), indicating that their coding capacity has been conserved during vertebrate evolution. One of these - a chuvirus-derived L-protein identified in livebearers – adds to previous evidence that viral RdRp sequences have been co-opted by vertebrate genomes [43]. Mapping of EVE loci can also inform efforts to develop new medical treatments - in a recent study, EVE loci identified using DIGS were used to identify potential genomic safe harbours for human transgene therapy applications [84].

The EVE screen performed here has several important limitations. Firstly, it relied on published WGS data generated for extant species. Secondly, our results have likely been influenced by aspects of our screening configuration, such as the composition of the probe set with respect to viral taxa and polypeptide probe length [85, 86]. This might mean that we failed to detect some of the potentially recognisable EVE loci present in our TDb. For example, counts of RT-encoding ERV loci were found to be generally lower in ray-finned fish and jawless fish (**Fig. 3**), but previous studies have shown that RT loci related to other families of reverse-transcribing virus, such as metaviruses (family *Metaviridae*) [87] and ‘lokiretroviruses’ [88] are relatively common in these hosts. These would likely have been missed in our search because they were not included in our RT RSL. Finally, previous studies have indicated that vertebrate genomes contain EVEs that lack any clear homology to extant viruses [89], and these would not be detected using a sequence similarity-based approach.

As vertebrate genome sequencing progresses, further opportunities to identify novel EVEs will arise, since: (i) any novel genome could in theory contain a lineage-specific EVE, and; (ii) ongoing characterization of exogenous virus diversity may allow for detection of previously undetectable EVEs. The DIGS project created here, which is openly available online, can be reused to accommodate newly sequenced vertebrate genomes (TDb expansion) and newly discovered vertebrate virus diversity (RSL/probe set expansion). In addition, similar projects can readily be created to screen for EVEs in other host groups.

The use of DIGS is not limited to investigations of EVEs. DIGS can be used to investigate any sufficiently conserved genome feature lurking within the dark genome, including both coding and non-coding elements (**Table 1**). Many of the most interesting genes have evolved relatively rapidly and are difficult to annotate reliably using automated approaches [90]. Furthermore, even relatively conserved genes may be incompletely annotated by automated pipelines. DIGS has previously been used to broadly survey the distribution of interferon stimulated genes in mammals [ref], and for in-depth investigation of specific genes and gene families, such as OAS1 [91] and APOBEC3 [92]. While DIGS is best suited to investigations of genome features that comprise a single contiguous unit and contain relatively long, easily recognised regions, it can also be used to investigate genome features that are shorter or are comprised of several short sub-components, providing that a careful approach is used. For example, when investigating interferon lambda (IFNL) genes, which are expressed from multiple, short exons, we included conserved flanking features in our RSL and probe set [93]. This enabled more confident matching of IFNL exons based on their positional relationships relative to conserved markers. We have also used DIGS in functional genomics studies to investigate the locations of short nucleotide motifs identified in binding assays (e.g. CHiP-seq) relative to other genomic features such as ERVs [94, 95].

The framework described here or implementing DIGS could be further developed and improved. For example, by including the option to use of other sequence similarity search tools, such as Diamond [96] and ElasticBLAST [97], and RNA structure based search tools such as INFERNAL [98]. Integrating with functional genomics resources could provide further dimensionality to the kinds of investigations that may be performed using DIGS [99].

## CONCLUSIONS

We demonstrate how a relational database management system can be linked to a similarity search-based screening pipeline to investigate the dark genome *in silico*. Using this approach, we catalogue and analyse EVEs throughout vertebrate genomes, providing a broad range of novel insights into the evolution of ancient viruses and their interactions with host species.

## MATERIALS & METHODS

### Whole genome sequence and taxonomic data

Whole genome shotgun (WGS) sequence assemblies of 874 vertebrate species were obtained from the NCBI genomes resource [100]. Taxonomic data for the vertebrate species included in our screen and the viruses in our reference sequence library were obtained from the NCBI taxonomy database [101], using PERL scripts included with the DIGS tool package.

### Database-integrated screening for RT-encoding ERVs

An RT RSL was collated to represent diversity within the *Retroviridae*. We included representatives of previously identified ERV lineages and exogenous retrovirus species. A subset of these sequences was used as probes in similarity search-based screens [42]. For initial screening we used a bitscore cut-off of 60. For comparisons of ERV RT copy number across species we filtered initial results using a more conservative bitscore cut-off of 90. Our previous, DIGS-based studies of ERVs have shown that spurious matches (i.e. to sequences other than retroviral RTs) do not arise when this cut-off is applied, although some genuine ERV RT hits may be excluded [67].

### Database-integrated screening for non-retroviral EVEs

We obtained an RSL representing the proteome of eukaryotic viruses from the NCBI virus genomes database [39]. We supplemented this with sequence likely to cross-match to virus probes during screening. These included the teratorn transposon found in fish, which is known to contain multiple alloherpesvirus-derived genes [102]. We included the polypeptide sequences of these genes, obtained from the subtype 1 Teratorn reference (Accession #: LC199500) in our RSL. We also included representatives of the maverick/polinton lineage of transposons, derived from sequences defined in a prevous study [103]. Since these element derive from a group of midsize eukaryotic linear dsDNA (MELD) viruses provisionally named ‘adintoviruses’ [71]. Probes constituted a subset of 685 sequences contained within our

RSL, and incorporated polypeptide sequences representing all major protein-coding genes of representative species of all recognised vertebrate virus families. We also included representative sequences of maverick/polinton elements in our probe set. We used a bit score cut-off of 60 as a threshold for counting non-retroviral EVE loci. This threshold was established through previous experience searching for non-retroviral EVEs using DIGS [48, 50, 51, 61]. Experience from previous studies had shown that nearly 100% of matches with bit scores >= 60 were either virus-derived or represented genuine similarity between virus genes and their cellular orthologs. By contrast, investigation of a subset of 100 hits with bit scores of b 40-59 showed that ∼50% could not be confidently confirmed as having a viral origin (data not shown).

Artefactual hits to host DNA can occur since some virus genomes contain genes that have cellular homologs [104], some virus genomes contain captured host DNA [105]. To distinguish host from virus-derived DNA in these cases, we exported such hits from the screening database and virtually translated them to obtain a polypeptide sequence. We then used the translated sequences as query input to online BLAST searches of GenBank’s non- redundant (nr) database. If searches revealed closer matching to host genes than to known viral genes, the input sequences were assumed to be host derived. Wherever this occurred we incorporated representatives of the matching host sequences into the RSL, so that they would be recognised as host hits on reclassification. By updating hit classifications in this way, we could progressively filter out host-derived hits from our final screening output.

### Filtering sequences-derived from exogenous viruses

Sequences derived from exogenous viruses are occasionally incorporated into WGS assemblies. We used SQL queries to identify and exclude these sequences based on hit characteristics. Where hits derived from virus species or species groups that have been sequenced previously, they could be discriminated on the basis of sequence identity (i.e., 98-99% nucleotide-level identity known viruses. The ‘extract start’ field could be used to identify sequences that lacked flanking genomic sequences, indicating a potential exogenous origin. We also examined the virtually translated sequences to look for evidence of long-term presence in the host germline (e.g., stop codons, frameshifting mutations).

### Filtering of cross-matching retrovirus-derived sequences

Hits that match more closely to virus genomes than to host DNA, and are clearly inserted into host DNA, are most likely *bona fide* EVE sequences. However, they may not necessarily be non-retroviral EVEs, because some filoviruses and arenaviruses (family *Arenaviridae*) contain glycoprotein genes that are distantly related to those found in certain retroviruses [106, 107]. When such hits were investigated and found to correspond to ERVs (established through the presence of proviral genome features adjacent to the hit) we included the putative sequences of glycoproteins encoded by these ERVs into our RSL and reclassified hits, so that spurious matches could be recognized as ERV-derived.

### Genomic analysis

Previous studies of presence/absence patterns have shown that non-retroviral EVEs are present in many genomes due to orthology (ancient insertions) rather than paralogy (recent independent insertion) [48, 50–52]. To differentiate orthologs of previously described EVEs from newly identified paralogs, we substituted expanded our RSL to include consensus/reference sequences representing unique EVE loci. This set of EVE loci was comprised of insertions identified in previous studies [48, 50, 51, 53, 108], as well as a set of clearly novel EVEs identified in the present screen. For high-copy number, amplified lineages within this set, we only included a single reference sequence, rather than attempting to represent each individual ortholog, since it was clear that these elements derive from a single germline incorporation event (see **Fig. S8**). EVEs were considered novel if: (i) they derived from a virus group not previously reported in the host group in which they were identified, or (ii) occurred in species only distantly related to species in which similar EVEs had been identified previously (e.g. an entirely distinct host class). Whenever novel EVEs were defined, results were reclassified using the updated RSL (see **Fig. 2**). Orthologs of previously identified EVEs could be inferred by using SQL queries to summarise screening results, as they disclosed high similarity to these EVE sequences and occurred in host species relatively closely related to the species in which the putatively orthologous EVEs had previously been identified. By contrast, novel paralogs either disclosed only limited similarity to previously identified EVE sequences or occurred in distantly related host species. This approach to discriminating between paralogs and orthologs has limitations, but can guide further investigations that use more reliable approaches (e.g. via investigation of flanking sequences, or phylogeny) to infer orthology [51]. Se-Al (version 2.0a11) was used to inspect multiple sequence alignments of EVEs and genomic flanking sequences. Minimum age estimates were obtained for orthologous EVEs by using host species divergence time estimates collated in TimeTree [109]. We identified open reading frames and open coding regions within EVEs using PERL scripts available on request.

### Phylogenetic analysis

Phylogenies were reconstructed using the maximum likelihood approach implemented in RAxML (version 8.2.12) [110] and model parameters selected using IQ-TREE model selection function [111]. Support for phylogenies was assessed via 1000 non-parametric bootstrap replicates. A time-calibrated vertebrate phylogeny was obtained via TimeTree, an open database of species divergence time estimates [109]. To determine germline infiltration rate, we divided the total number of distinct EVE orthologs identified in each vertebrate class by the total amount of branch length sampled for that class (obtained from the time- calibrated phylogeny).

### Application of standardised nomenclature to EVE loci

We assigned all non-retroviral EVEs identified in our study unique identifiers (IDs), following a convention developed for ERVs [112], Each was assigned a unique identifier (ID) constructed from three components. The first component is a classifier denoting the type of EVE. The second component comprises: (i) the name of the taxonomic group of viruses the element derived from and; (ii) a numeric ID that uniquely identifies a specific integration locus, or for multicopy lineages, a unique founding event. The final component denotes the taxonomic distribution of the element. This approach has been applied in several previous studies of vertebrate EVEs [50, 51, 53, 61] and we maintained consistency with these studies with respect to the numeric ID. Where our study revealed new information about the taxonomic relationship of an EVE to contemporary viruses, or its distribution across taxa, the ID was updated accordingly.

## DECLARATIONS

### Ethics approval and consent to participate

Not applicable

### Consent for publication

Not applicable

### Availability of data and materials

All data are openly available in the DIGS-for-EVEs repository hosted on GitHub: https://github.com/giffordlabcvr/DIGS-for-EVEs

https://twitter.com/DigsTool

### Competing interests

None declared.

### Funding

DB-M is supported by the V Foundation for Cancer Research and the Searle Scholars Program. TD, SL, JH, and RJG, were funded by the Medical Research Council of the United Kingdom (MC_UU_12014/12).

## Acknowledgments

We thank Connor Bamford, Paul Bieniasz, and Jamie Henzy for helpful discussions. Additional thanks to Ade Tukuru (Aaron Diamond AIDS Research Centre) and Scott Arkinson (MRC-University of Glasgow Centre for Virus Research) for bioinformatics support.

## Notes

https://github.com/giffordlabcvr/DIGS-for-EVEs

### Competing Interest Statement

The authors have declared no competing interest.

### Summary of Updates

This version of the manuscript incorporates some minor edits and an extension to supplementary figures. Large supplementary tables have been removed from the manuscript and are instead available in a manuscript-associated public repository, referenced within the manuscript.

