## Supplementary material for "A novel approach to exploring the dark genome and its application to mapping of the vertebrate virus ‘fossil record’": Table S1

**Table S1. Putatively exogenous viruses identified in WGS data**

| **Vertebrate species^a^** | **Virus family/genus** | **Contig** |
| --- | --- | --- |
| **dsDNA** |  |  |
| Bumblebee bat (*Craseonycteris thonglongyai* | *Adenoviridae** | PVKE010106839.1, PVKE010087958.1 |
| Gracile opossum (*Gracilinanus agilis)* | *Adenoviridae** | JADWME010000011.1 |
| Viscacha rat (*Octomys mimax*) |  | NDGM010762229.1, NDGM010239277.1, NDGM010144971.1, NDGM010804184.1 |
| Ord's kangaroo rat (*Dipodomys ordii)* | *Adenoviridae** | scaffold_25807_dna |
| Greater bamboo lemur (*Prolemur simus*) | *Papillomaviridae** | MPIZ01127437.1 |
| Humpbacked dolphin (*Sousa chinensis*) | *Papillomaviridae** | QWLN01062956.1, QWLN01040988.1 |
| Duck-billed platypus (*Ornithorhynchus anatinus*) | *Papillomaviridae* | Contig159295 |
| Budgerigar (*Melopsittacus undulatus*) | *Herpesviridae* | JH541059 |
| **ssDNA** |  |  |
| David's myotis (*Myotis davidii*) | *Chaphamaparvovirus* | KB106247.1 |
| White-faced capuchin (*Cebus imitator*) | *Chaphamaparvovirus* | KV391748.1 |
| Brown mesite (*Mesitornis unicolor*) | *Chaphamaparvovirus* | Scaffold28758 |
| Pit viper (*Protobothrops mucrosquamatus*) | *Chaphamaparvovirus* |  |
| David's myotis (*Myotis davidii*) | *Chaphamaparvovirus* |  |
| Canary (*Serinus canaria*) | *Chaphamaparvovirus* |  |
| Gulf pipefish | *Ichthamaparvovirus* |  |
| Senegalese bichir | *Protoparvovirus* |  |
| **ssRNA+ve** |  |  |
| Vicugna pacos | *Hepacivirus* | ABRR02259018 |

**Legend:** **^a^** Common name (Latin binomial). * Previously unreported, putatively novel virus species.
