## Supplementary Figure Legends for "A novel approach to exploring the dark genome and its application to mapping of the vertebrate virus ‘fossil record’"

**Figure S1. An annotated example of a DIGS tool control file**.

The DIGS tool control file defines parameters for screening and the paths to reference sequence library (RSL), probe, and target database (TDb) files. Control files are structured as ‘NEXUS’ style blocks [1] delineated by BEGIN and ENDBLOCK tokens.

**Figure S2. The DIGS tool framework for *in silico* genome screening.**

Panels show schematic representations of key components of the *in silico* genome screening framework that is implemented in the database-integrated (DIGS) tool. Colours indicate relationships between components.

***(a) Screening database schema*.**

The database schema consists of three essential tables: (1) a status table ('searches performed'), (2) a data processing table ('active set'), and (3) a results table ('digs results').

These tables serve distinct purposes within the screening process:

*Status Table:* This table is used to monitor progress by keeping a record of the searches performed. It tracks progress in screening, by recording BLAST-based queries (i.e. which probes have so far been used to screen which TDb files).

*Data Processing Table:* The data processing table is employed by the DIGS screening algorithm. It temporarily stores and analyses the outcomes of individual BLAST searches and is cleared after each screening iteration.

*Results Table:* The results table holds a unique collection of all the hits obtained during the screening. It includes fields that define: (i) the location and orientation of a contiguous region of sequence within a specific genomic scaffold within a specific TDb file; (ii) the closest RSL match to each hit, as determined by BLAST comparison (see **Fig. 2** in main text), and associated values.

Additional flexibility is provided by the ability to expand DIGS screening databases with extra tables linked to the core schema using designated fields, as indicated in the figure by coloured circles. The inset box offers two examples of custom tables utilized in this study:

*Virus Taxonomy Table*: This table is linked via the 'assigned name' field.

*Host Taxonomy Table*: This table is linked via the 'organism' field.

***(b) Target database directory hierarchy.*** The DIGS tool requires that TDb files are arranged in a directory hierarchy as shown. Directory levels correspond to screening database fields as shown in panel (a). Concatenating the values of four fields shown in darker blue (i.e. ‘organism’, ‘data type’, and ‘assembly version’, and ‘target name’) forms a unique identifier for target files. These fields define, respectively, the species of genome data origin, a user-defined data ‘type’ (e.g., ‘sanger genome’, ‘ngs-transcriptome’), assembly version name, and a specific genome assembly file. Extending this concatenation to include the ‘scaffold’, ‘extract start’ and ‘extract end’ fields defines a locus within the TDb.

***(c) Reference sequence library.*** The reference sequence library (RSL) is a representative set of reference sequences for the genome feature(s) under investigation. FASTA headers encode two levels of classification; (i) the name of the species from which the sequence derives; (ii) the name of the genome feature represented by the sequence.

**Figure S3. Examples of SQL-based querying of DIGS results.**

Panels show screenshots of a MySQL client program (Sequel Ace) connected to a screening database generated using the database-integrated genome screening (DIGS) tool. Structured query language (SQL) can be used to interrogate and manipulate screening databases. Each panel shows a distinct SQL query (upper window) and its results (lower window).

**(a)** Deriving stratified counts of non-retroviral EVE loci. The query shown here uses a custom table (‘virus taxonomy’,) linked via the ‘assigned name’ field (see **Fig. S**, to group hits according to virus family. A ‘digs results’ table field, ‘bit score’, is used to restrict results to higher confidence hits - it is derived from the ‘reverse’ BLAST-step, in which a sequence hit identified from a ‘forward’ BLAST (i.e. probe versus target database file) is compared to the reference sequence library (RSL). It thereby provides an index showing how similar individual hits are to sequences included in the RSL.

**(b)** Retrieving information about paramyxovirus-derived EVEs recovered via DIGS. The query shown here provides an overview of paramyxovirus EVEs, showing the vertebrate class in which they were identified, which paramyxovirus species and genes they disclose similarity to (based on comparison to RSL sequences), and fields that show the closeness of the match. It uses two custom tables: (i) ‘virus taxonomy’ and (ii) ‘host taxonomy,’ which is linked via the ‘organism’ field (see **Fig. S2**). A bitscore cut-off is used to filter hits, and an order statement is used to sort results by the taxonomic designations of host species.

**(c)** An SQL query used to retrieve individual counts of high confidence endogenous retrovirus (ERV) reverse transcriptase (RT). RT hits were recovered by screening a target database comprising whole genome sequence data of 874 vertebrate species. We empirically determined that a bit score of >=90 eliminates false positive RT hits. Host and virus taxonomy tables, linked as described above for non-retroviral viruses, are used to stratify results.

**Figure S4. Validation of the DIGS tool.**

Tetherin ortholog data was downloaded from OrthoDB and Ensembl. Results from all three pipelines were compared based on the Ensembl Protein ID and were found to overlap by >99% as shown above.

.

**Figure S5.** **An unusual retrovirus-like sequence identified in the genome of the pig-nosed turtle.**

**(a)** Predicted ORFs in the region of the EVE locus, determined using NCBI’s ‘ORF finder’ program [2]. The sequence input to ORF-finder included 5 kilobases of DNA sequence flanking the EVE locus. **(b)** tBLASTn result showing similarity of largest ORF to equine foamy virus. **(c)** tBLASTn result showing similarity between ORF 60 and a chelonid (turtle) herpesvirus.

**Figure S6. Summary of vertebrate EVE coding potential.**

Histograms showing the distribution of lengths, binned in increments of 100 amino acid (aa) residues, of uninterrupted coding regions (lighter bars) and open reading frames (darker bars) longer than 300aa, contained within vertebrate endogenous viral elements (EVEs). was plotted in R using ggplot with ‘geom hist’ function.

**Figure S7. Evolutionary relationships of vertebrate EVEs and viruses.**

Bootstrapped maximum likelihood trees showing the reconstructed evolutionary relationships between vertebrate EVEs and related viruses. **(a)** Gammaherpesvirus *terminase*; **(b)** Alphaherpesvirus *glycoprotein B*; **(c)** Alloherpesvirus *terminase*; **(d)** Circovirus *rep*; **(e)** Parvovirus *rep*; **(f)** Hepadnavirus *pol*; **(g)** Paramyxovirus *L polymerase*; **(h)** Chuvirus *nucleoprotein*; **(i)** Bornavirus *L polymerase*; **(j)** Bornavirus *glycoprotein*. Numbers on nodes indicate bootstrap support (100 bootstrap replicates). All trees were constructed from nucleotide level alignments using the General Time Reversible (GTR) model of nucleotide substitution. Scale bars show evolutionary distance in substitutions per site.

**Figure S8. Amplified lineages of endogenous viral elements.**

**Panels (a-b)** Maximum likelihood (ML) phylogenies showing reconstructed evolutionary relationships among **(a)** an lineage of endogenous hepatitis B (eHBV) elements, labelled ‘ehbv-avi.27-Suliformes’, that has been amplified in cormorant (family Phalacrocoracidae) genomes and; **(b)** a lineage of long interspersed nuclear element (LINE)-associated endogenous circoviral elements (ECVs), labelled ‘ecv-circo.51-Carnivora’, amplified in carnivore genomes (order Carnivora). ML phylogenies were reconstructed from nucleotide-level alignments. Scale bars show evolutionary distance in substitutions per site. **(c)** Genomic structure found in an ecv-circo.51-Carnivora subclade, comprised of a LINE1 homologous region (left) and region homologous to a circoviral *rep* gene (right). The orientation of each of these sub-components is indicated, and the presence of a poly-adenine (A) tail. Homology of the LINE1 region to human gammaherpesvirus 4 and an isolate of Chikungunya virus is indicated in panel **(d)** and **(e)**, respectively.
