## Supplementary Figures for "A novel approach to exploring the dark genome and its application to mapping of the vertebrate virus ‘fossil record’"

```
# DIGS screening database connection parameters

Begin SCREENDB;
            db_name=eve_1_parvoviridae;
            mysql_server=localhost;
ENDBLOCK;


# Paths and parameters for in silico screening using DIGS

BEGIN SCREENSETS;
            query_aa_fasta=/parvo-probes.faa;
            reference_aa_fasta=/parvo-refs.faa;
               output_path=./tmp/;
            bitscore_min_tblastn=60;
            seq_length_minimum=40;
            defragment_range=10;
            consolidate_range=3000;
            blast_threads=8;
ENDBLOCK;


# List of target genomes to screen

BEGIN TARGETS;
            Agnatha/
            Actinopterygii/
            Sarcopterygii/
            Chondrichthyes/
            Amphibia/
            Squamata/
            Crocodilia/
            Aves/
            Mammalia/
ENDBLOCK;
```

*Screening DB connection details*

*— Path to  the query sequences in FASTA format*

*— Path to reference sequence library (RSL)*

*Params*

*Target database*

**Figure S1.**

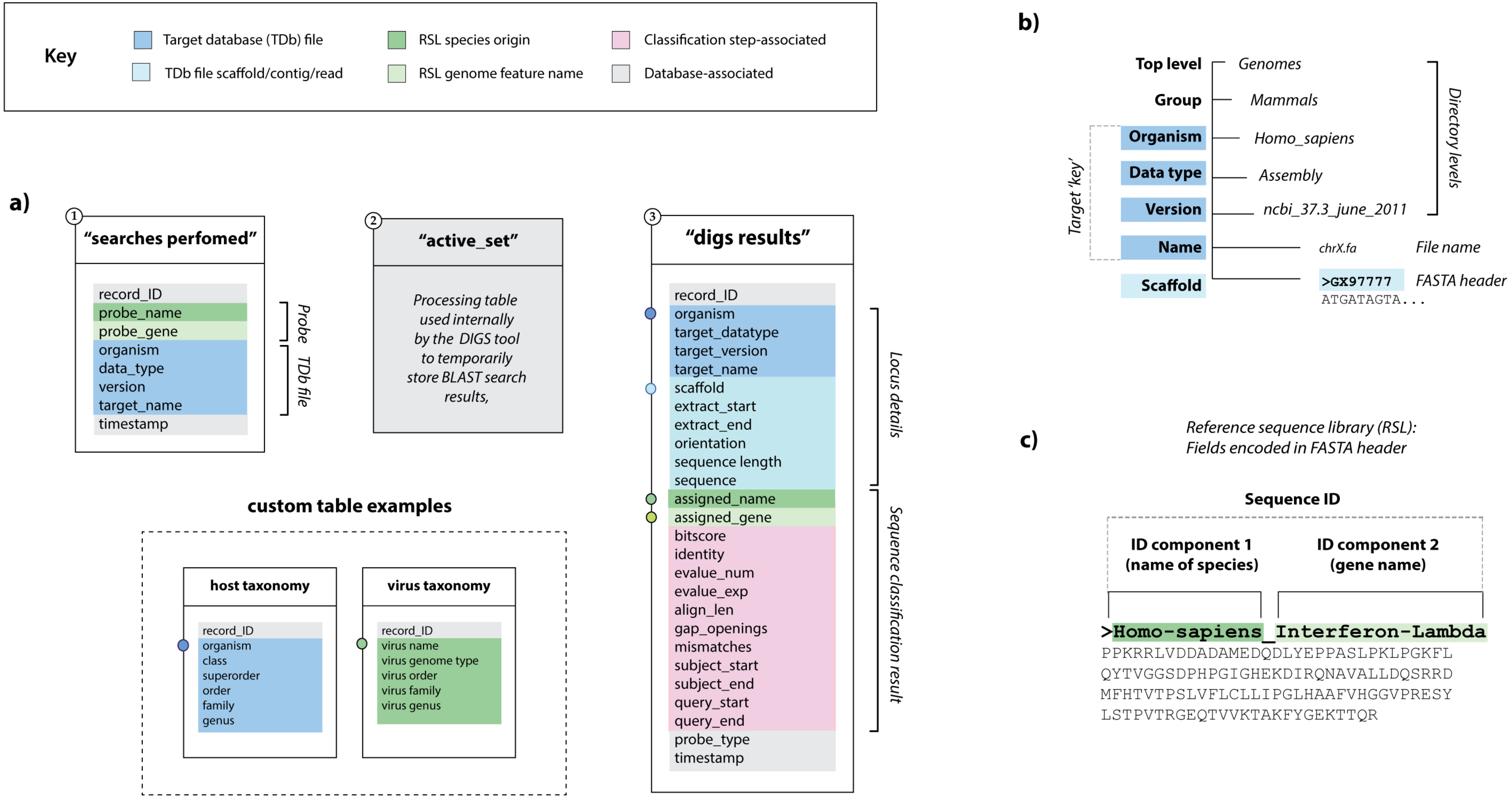

Key
Target database (TDb) file
TDb file scaffold/contig/read
RSL species origin
RSL genome feature name
Classification step-associated
Database-associated
a)
1
"searches perfomed"
record_ID
probe_name
probe_gene
organism
data_type
version
target_name
timestamp
Probe
TDb file
2
"active_set"
Processing table used internally by the DIGS tool to temporarily store BLAST search results,
3
"digs results"
record_ID
organism
target_datatype
target_version
target_name
scaffold
extract_start
extract_end
orientation
sequence length
sequence
assigned_name
assigned_gene
bitscore
identity
evalue_num
evalue_exp
align_len
gap_openings
mismatches
subject_start
subject_end
query_start
query_end
probe_type
timestamp
Locus details
Sequence classification result
custom table examples
host taxonomy
record_ID
organism
class
superorder
order
family
genus
virus taxonomy
record_ID
virus name
virus genome type
virus order
virus family
virus genus
b)
Top level
Genomes
Group
Mammals
Organism
Homo_sapiens
Data type
Assembly
Version
ncbi_37.3_june_2011
Name
chrX.fa
File name
Scaffold
>GX97777
ATGATAGTA...
FASTA header
Target 'key'
Directory levels
c)
Reference sequence library (RSL):
Fields encoded in FASTA header
Sequence ID
ID component 1
(name of species)
ID component 2
(gene name)
>Homo-sapiens_Interferon-Lambda
PPKRRLVDDADAMEDQDLYEPPASLPKLPGKFL
QYTVGGSDPHPGIGHEKDIRQNAVALLDQSRRD
MFHTVTPSLVFLCLLIPGLHAAFVHGGVPRESY
LSTPVTRGEQTVVKTAKFYGEKTTQR


**Figure S2.**

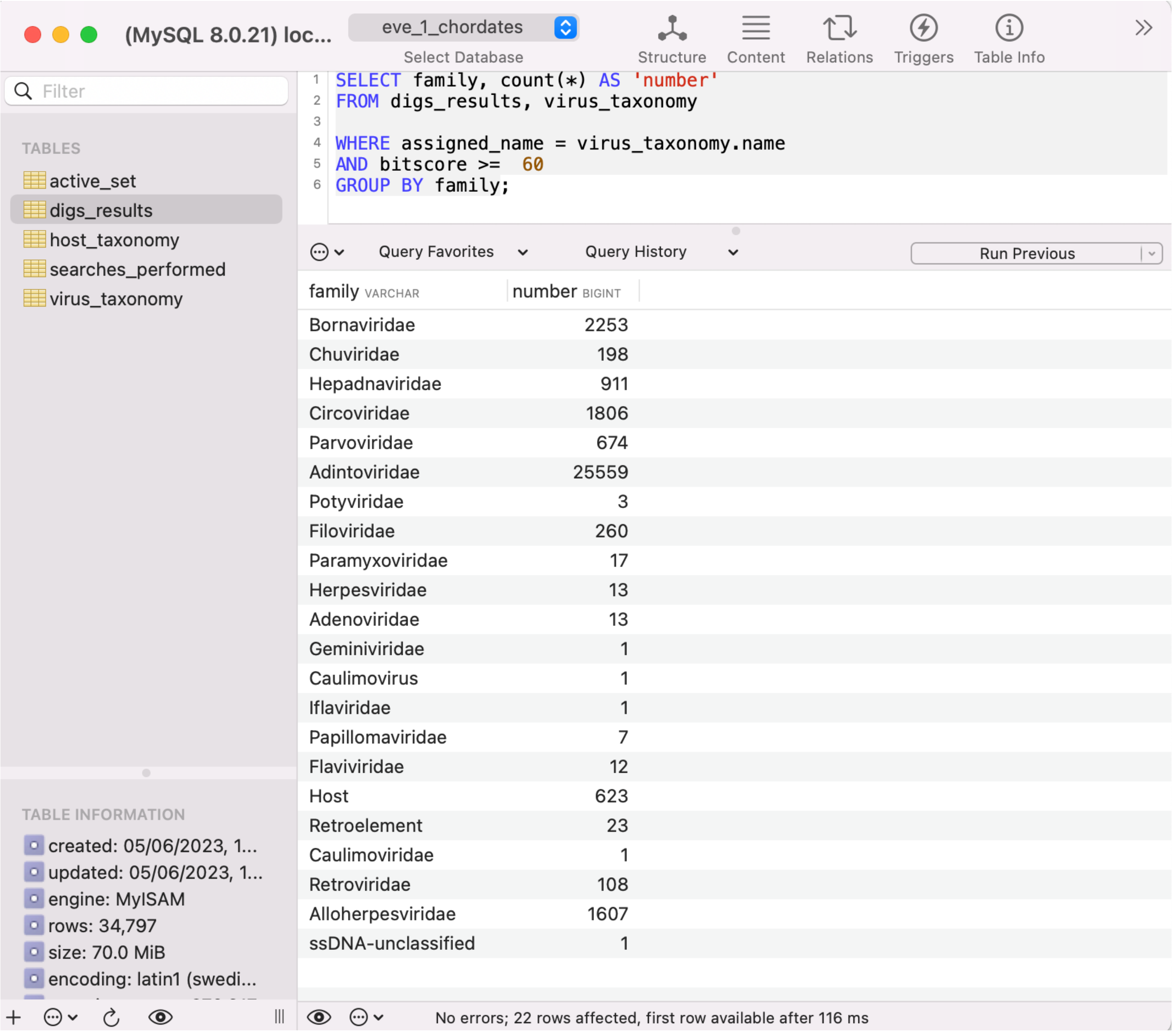
(MySQL 8.0.21) loc...
eve_1_chordates
Select Database
Structure
Content
Relations
Triggers
Table Info
Filter
TABLES
active_set
digs_results
host_taxonomy
searches_performed
virus_taxonomy
SELECT family, count(*) AS 'number'
FROM digs_results, virus_taxonomy
WHERE assigned_name = virus_taxonomy.name
AND bitscore >= 60
GROUP BY family;
Query Favorites
Query History
Run Previous
family VARCHAR
number BIGINT
Bornaviridae 2253
Chuviridae 198
Hepadnaviridae 911
Circoviridae 1806
Parvoviridae 674
Adintoviridae 25559
Potyviridae 3
Filoviridae 260
Paramyxoviridae 17
Herpesviridae 13
Adenoviridae 13
Geminiviridae 1
Caulimovirus 1
Iflaviridae 1
Papillomaviridae 7
Flaviviridae 12
Host 623
Retroelement 23
Caulimoviridae 1
Retroviridae 108
Alloherpesviridae 1607
ssDNA-unclassified 1
TABLE INFORMATION
created: 05/06/2023, 1...
updated: 05/06/2023, 1...
engine: MyISAM
rows: 34,797
size: 70.0 MiB
encoding: latin1 (swedi...
No errors; 22 rows affected, first row available after 116 ms

**Figure S3a.**

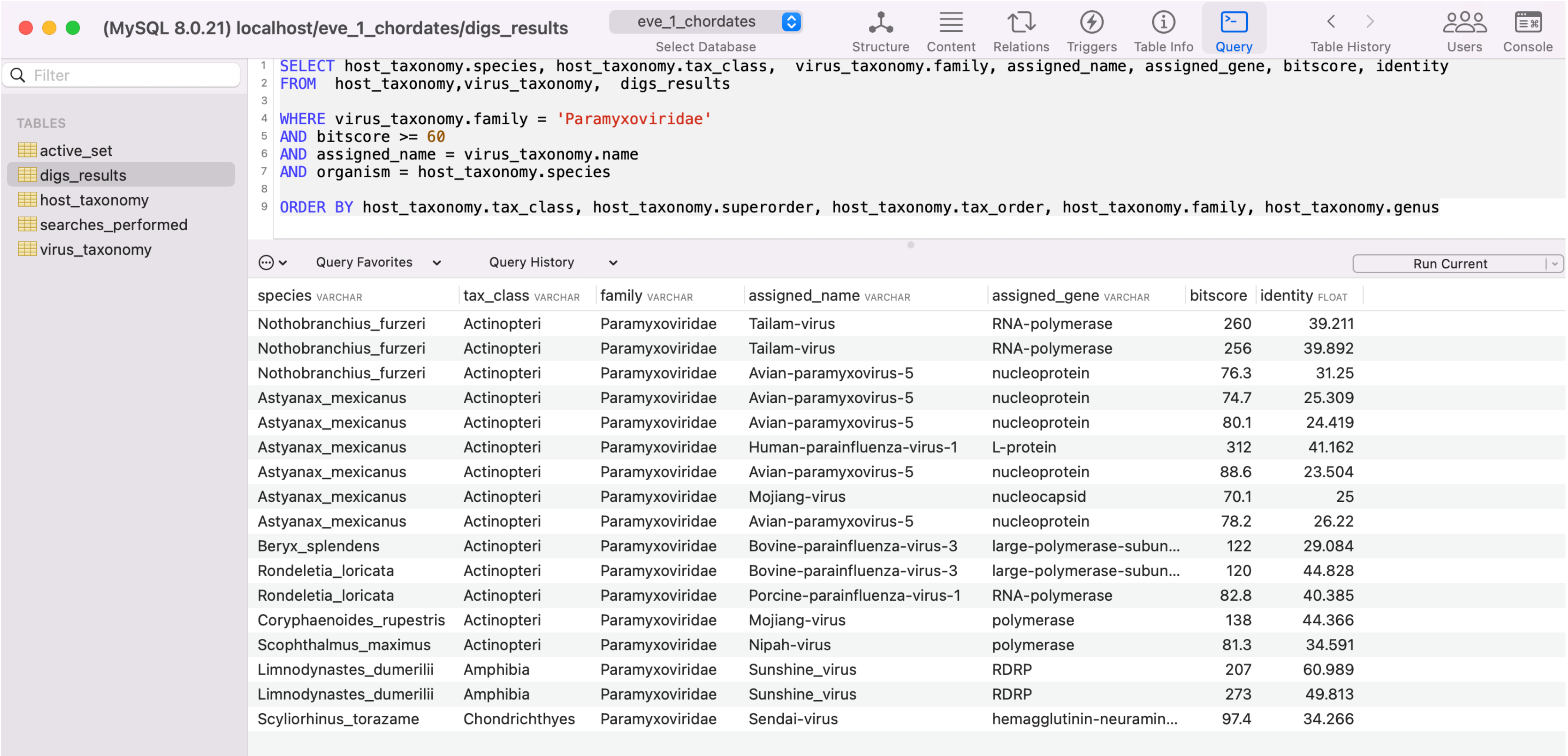


```
SELECT host_taxonomy.species, host_taxonomy.tax_class,  virus_taxonomy.family, assigned_name, assigned_gene, bitscore, identity
FROM  host_taxonomy,virus_taxonomy,  digs_results

WHERE virus_taxonomy.family = 'Paramyxoviridae'
AND bitscore >= 60
AND assigned_name = virus_taxonomy.name
AND organism = host_taxonomy.species

ORDER BY host_taxonomy.tax_class, host_taxonomy.superorder, host_taxonomy.tax_order, host_taxonomy.family, host_taxonomy.genus
```

| species VARCHAR | tax_class VARCHAR | family VARCHAR | assigned_name VARCHAR | assigned_gene VARCHAR | bitscore | identity FLOAT |
|---|---|---|---|---|---|---|
| Nothobranchius_furzeri | Actinopteri | Paramyxoviridae | Tailam-virus | RNA-polymerase | 260 | 39.211 |
| Nothobranchius_furzeri | Actinopteri | Paramyxoviridae | Tailam-virus | RNA-polymerase | 256 | 39.892 |
| Nothobranchius_furzeri | Actinopteri | Paramyxoviridae | Avian-paramyxovirus-5 | nucleoprotein | 76.3 | 31.25 |
| Astyanax_mexicanus | Actinopteri | Paramyxoviridae | Avian-paramyxovirus-5 | nucleoprotein | 74.7 | 25.309 |
| Astyanax_mexicanus | Actinopteri | Paramyxoviridae | Avian-paramyxovirus-5 | nucleoprotein | 80.1 | 24.419 |
| Astyanax_mexicanus | Actinopteri | Paramyxoviridae | Human-parainfluenza-virus-1 | L-protein | 312 | 41.162 |
| Astyanax_mexicanus | Actinopteri | Paramyxoviridae | Avian-paramyxovirus-5 | nucleoprotein | 88.6 | 23.504 |
| Astyanax_mexicanus | Actinopteri | Paramyxoviridae | Mojiang-virus | nucleocapsid | 70.1 | 25 |
| Astyanax_mexicanus | Actinopteri | Paramyxoviridae | Avian-paramyxovirus-5 | nucleoprotein | 78.2 | 26.22 |
| Beryx_splendens | Actinopteri | Paramyxoviridae | Bovine-parainfluenza-virus-3 | large-polymerase-subun... | 122 | 29.084 |
| Rondeletia_loricata | Actinopteri | Paramyxoviridae | Bovine-parainfluenza-virus-3 | large-polymerase-subun... | 120 | 44.828 |
| Rondeletia_loricata | Actinopteri | Paramyxoviridae | Porcine-parainfluenza-virus-1 | RNA-polymerase | 82.8 | 40.385 |
| Coryphaenoides_rupestris | Actinopteri | Paramyxoviridae | Mojiang-virus | polymerase | 138 | 44.366 |
| Scophthalmus_maximus | Actinopteri | Paramyxoviridae | Nipah-virus | polymerase | 81.3 | 34.591 |
| Limnodynastes_dumerilii | Amphibia | Paramyxoviridae | Sunshine_virus | RDRP | 207 | 60.989 |
| Limnodynastes_dumerilii | Amphibia | Paramyxoviridae | Sunshine_virus | RDRP | 273 | 49.813 |
| Scyliorhinus_torazame | Chondrichthyes | Paramyxoviridae | Sendai-virus | hemagglutinin-neuramin... | 97.4 | 34.266 |

**Figure S3b.**

(MySQL 8.0.21) localhost/erv_0_rt_vertebrates

erv_0_rt_vertebrates
Select Database
Structure

Filter

TABLES

- active_set
- blast_chains
- digs_results
- host_taxonomy
- loci
- loci_chains
- rv_taxonomy
- searches_performed

```sql
SELECT  host_taxonomy.host_class, rv_taxonomy.clade, COUNT(*) as 'Number'

FROM digs_results, rv_taxonomy, host_taxonomy

WHERE digs_results.assigned_name = rv_taxonomy.name
AND host_taxonomy.species = organism
AND bitscore >= 90


GROUP BY host_taxonomy.host_class, rv_taxonomy.clade

ORDER BY host_taxonomy.host_class, rv_taxonomy.clade
```

Query Favorites    Query History

| host_class VARCHAR | clade VARCHAR | Number BIGINT |
|---|---|---|
| Actinistia | III | 97 |
| Actinopteri | I | 8514 |
| Actinopteri | II | 64 |
| Actinopteri | III | 2177 |
| Agnatha | I | 32 |
| Agnatha | II | 1 |
| Agnatha | III | 300 |
| Amphibia | I | 17319 |
| Amphibia | II | 973 |
| Amphibia | III | 8019 |
| Aves | I | 17951 |
| Aves | II | 20797 |
| Aves | III | 42014 |
| Chondrichthyes | I | 2018 |
| Chondrichthyes | III | 2843 |
| Mammalia | I | 215304 |
| Mammalia | II | 174549 |
| Mammalia | III | 143364 |
| Reptilia | I | 13676 |
| Reptilia | II | 12120 |
| Reptilia | III | 20197 |

**Figure S3c**

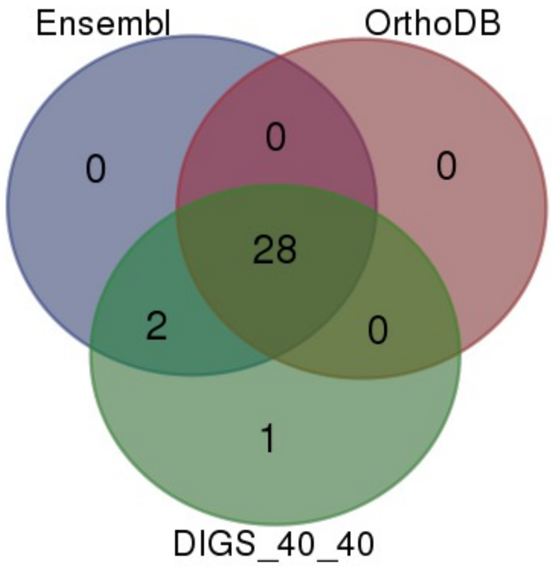


**Figure S4.**

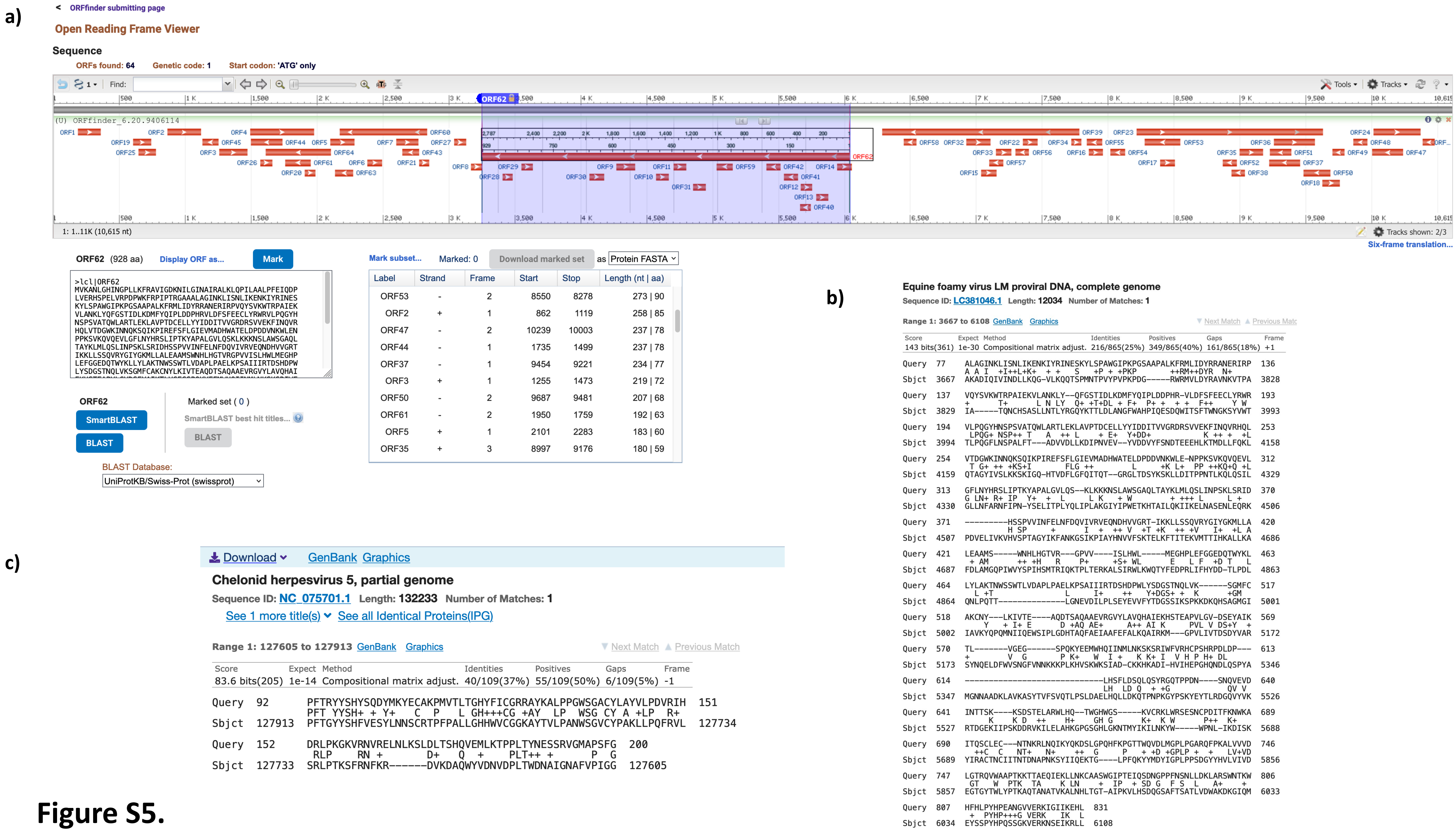
a)
ORFfinder submitting page
Open Reading Frame Viewer
Sequence
ORFs found: 64 Genetic code: 1 Start codon: 'ATG' only
1: 1..11K (10,615 nt)
Tracks shown: 2/3
Six-frame translation...
ORF62 (928 aa) Display ORF as... Mark
Mark subset... Marked: 0 Download marked set as Protein FASTA
Label Strand Frame Start Stop Length (nt | aa)
ORF53 - 2 8550 8278 273 | 90
ORF2 + 1 862 1119 258 | 85
ORF47 - 2 10239 10003 237 | 78
ORF44 - 1 1735 1499 237 | 78
ORF37 - 1 9454 9221 234 | 77
ORF3 + 1 1255 1473 219 | 72
ORF50 - 2 9687 9481 207 | 68
ORF61 - 2 1950 1759 192 | 63
ORF5 + 1 2101 2283 183 | 60
ORF35 + 3 8997 9176 180 | 59
ORF62
SmartBLAST
BLAST
Marked set ( 0 )
SmartBLAST best hit titles...
BLAST
BLAST Database:
UniProtKB/Swiss-Prot (swissprot)
b)
Equine foamy virus LM proviral DNA, complete genome
Sequence ID: LC381046.1 Length: 12034 Number of Matches: 1
Range 1: 3667 to 6108 GenBank Graphics
Score Expect Method Identities Positives Gaps Frame
143 bits(361) 1e-30 Compositional matrix adjust. 216/865(25%) 349/865(40%) 161/865(18%) +1
c)
Download GenBank Graphics
Chelonid herpesvirus 5, partial genome
Sequence ID: NC_075701.1 Length: 132233 Number of Matches: 1
See 1 more title(s) See all Identical Proteins(IPG)
Range 1: 127605 to 127913 GenBank Graphics
Next Match Previous Match
Score Expect Method Identities Positives Gaps Frame
83.6 bits(205) 1e-14 Compositional matrix adjust. 40/109(37%) 55/109(50%) 6/109(5%) -1
Query 92 PFTRYYSHYSQDYMKYECAKPMVTLTGHYFICGRRAYKALPPGWSGACYLAYVLPDVRIH 151
Sbjct 127913 PFTGYYSHFVESYLNNSCRTPFPALLGHHWVCGGKAYTVLPANWSGVCYPAKLLPQFRVL 127734
Query 152 DRLPKGKVRNVRELNLKSLDLTSHQVEMLKTPPLTYNESSRVGMAPSFG 200
Sbjct 127733 SRLPTKSFRNFKR-------DVKDAQWYVDNVDPLTWDNAIGNAFVPIGG 127605

### Figure S5.

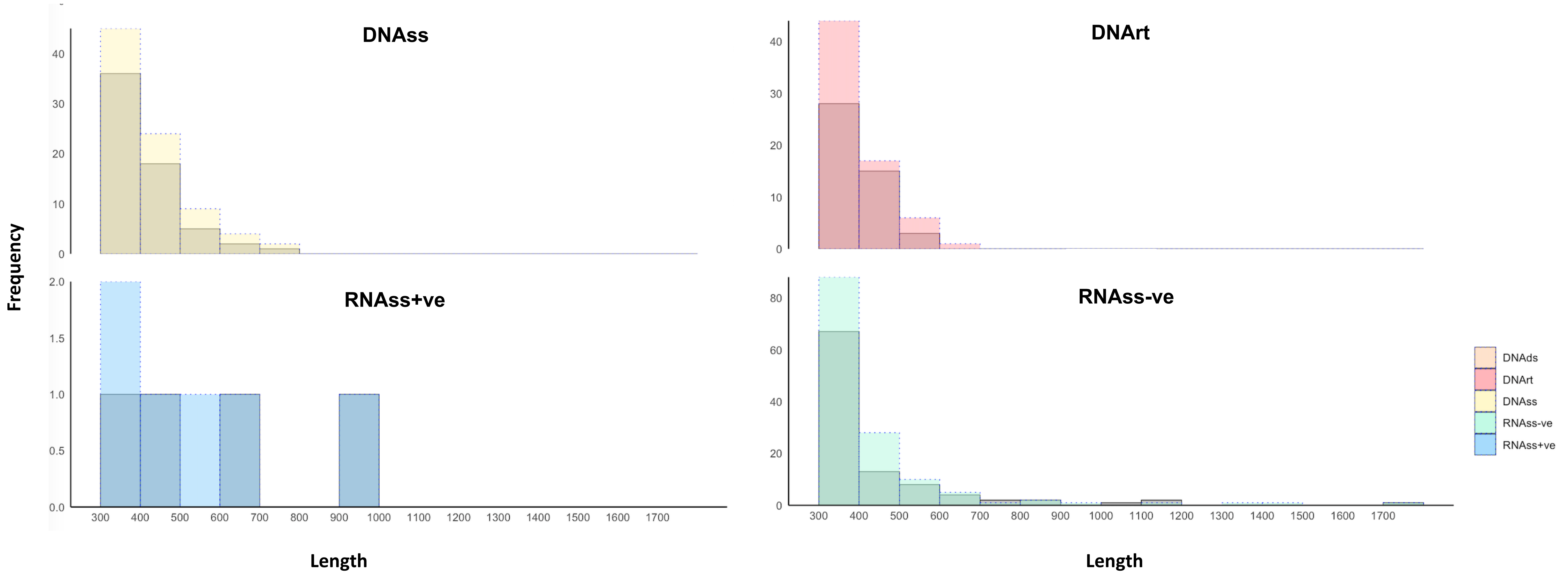
DNAss
DNArt
RNAss+ve
RNAss-ve
Frequency
Length
Length
0
10
20
30
40
0.0
0.5
1.0
1.5
2.0
60
80
300
400
500
600
700
800
900
1000
1100
1200
1300
1400
1500
1600
1700
DNAds
DNArt
DNAss
RNAss-ve
RNAss+ve

**Figure S6.**

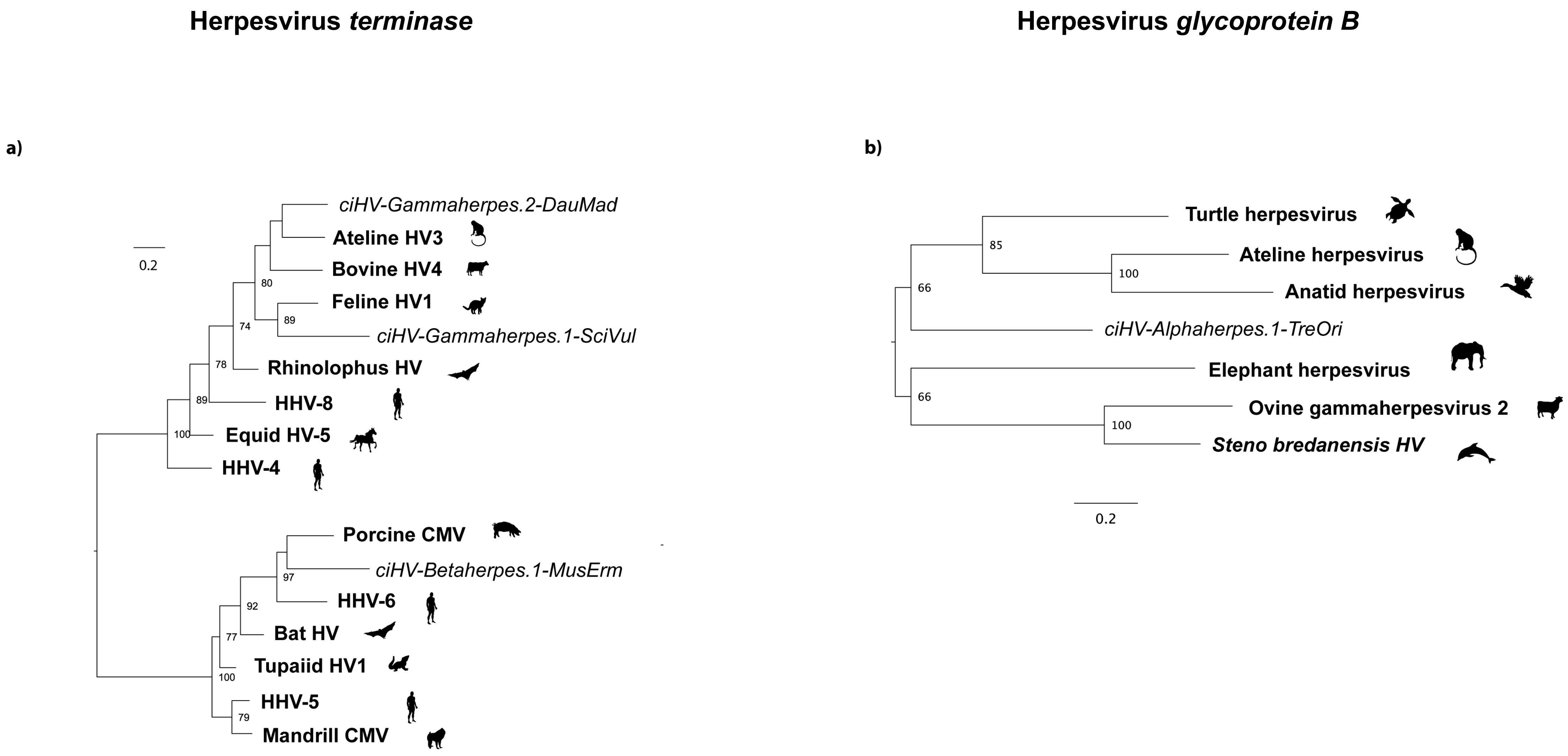
Herpesvirus terminase
a)
0.2
ciHV-Gammaherpes.2-DauMad
Ateline HV3
Bovine HV4
80
Feline HV1
89
74
ciHV-Gammaherpes.1-SciVul
78
Rhinolophus HV
89
HHV-8
100
Equid HV-5
HHV-4
Porcine CMV
ciHV-Betaherpes.1-MusErm
97
92
HHV-6
77
Bat HV
Tupaiid HV1
100
HHV-5
79
Mandrill CMV
Herpesvirus glycoprotein B
b)
Turtle herpesvirus
85
Ateline herpesvirus
100
Anatid herpesvirus
66
ciHV-Alphaherpes.1-TreOri
Elephant herpesvirus
66
Ovine gammaherpesvirus 2
100
Steno bredanensis HV
0.2

**Figure S7a-b.**

#### Alloherpesvirus *terminase*

#### Circovirus *rep*

c)

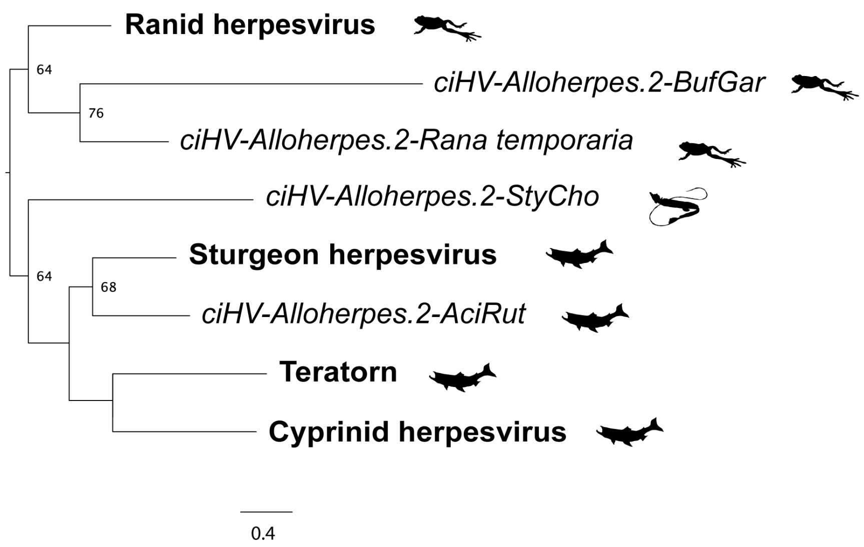


d)

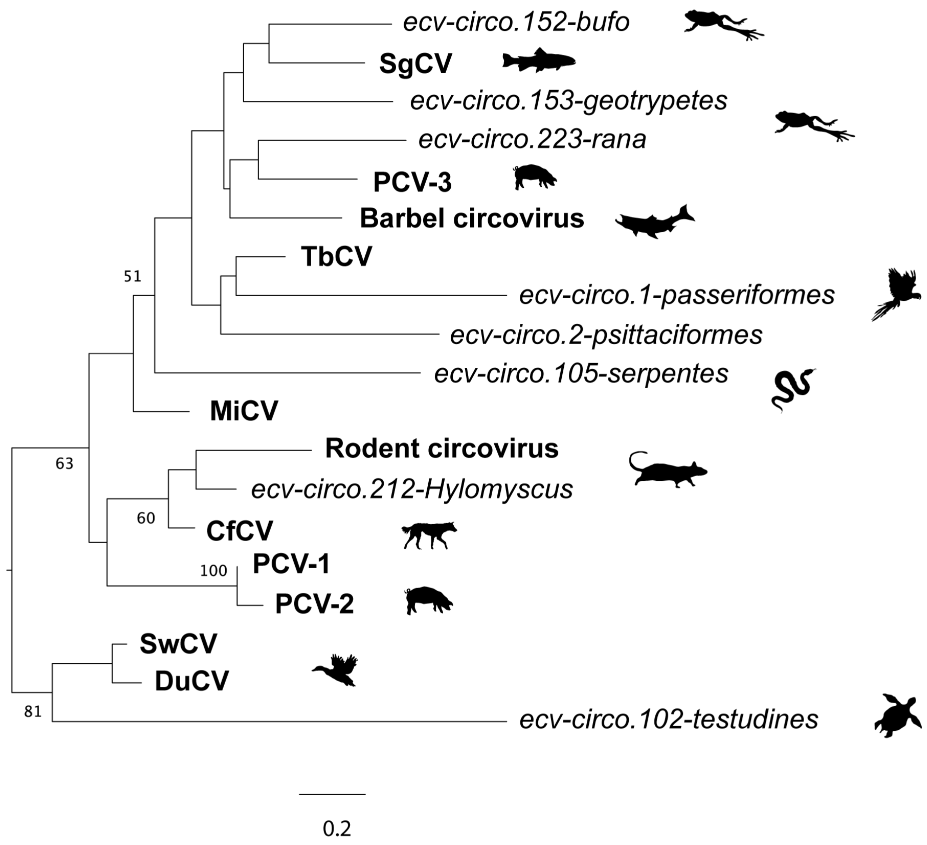


**Figure S7c-d.**

#### Parvovirus *rep*

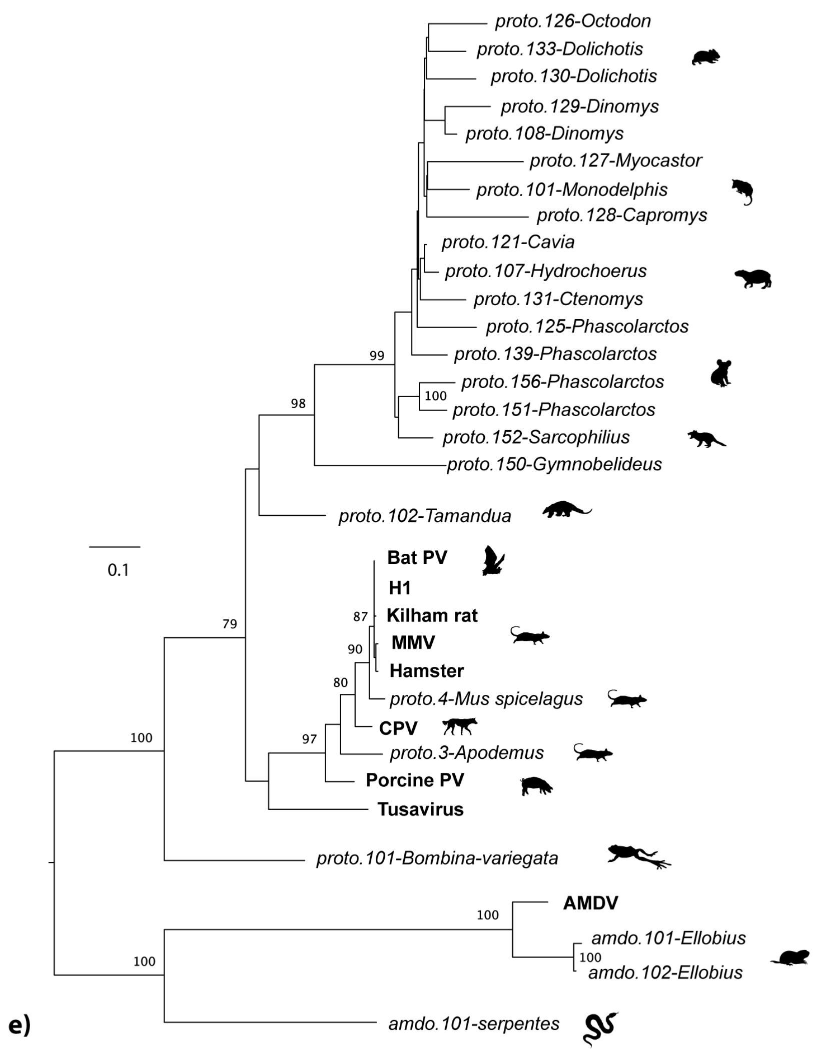


#### Hepadnavirus *pol*

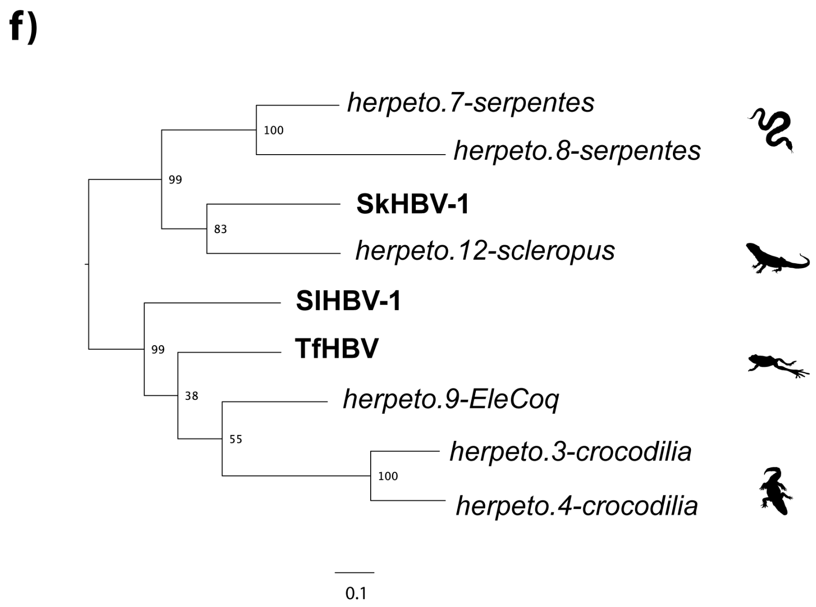


**Figure S7e-f.**

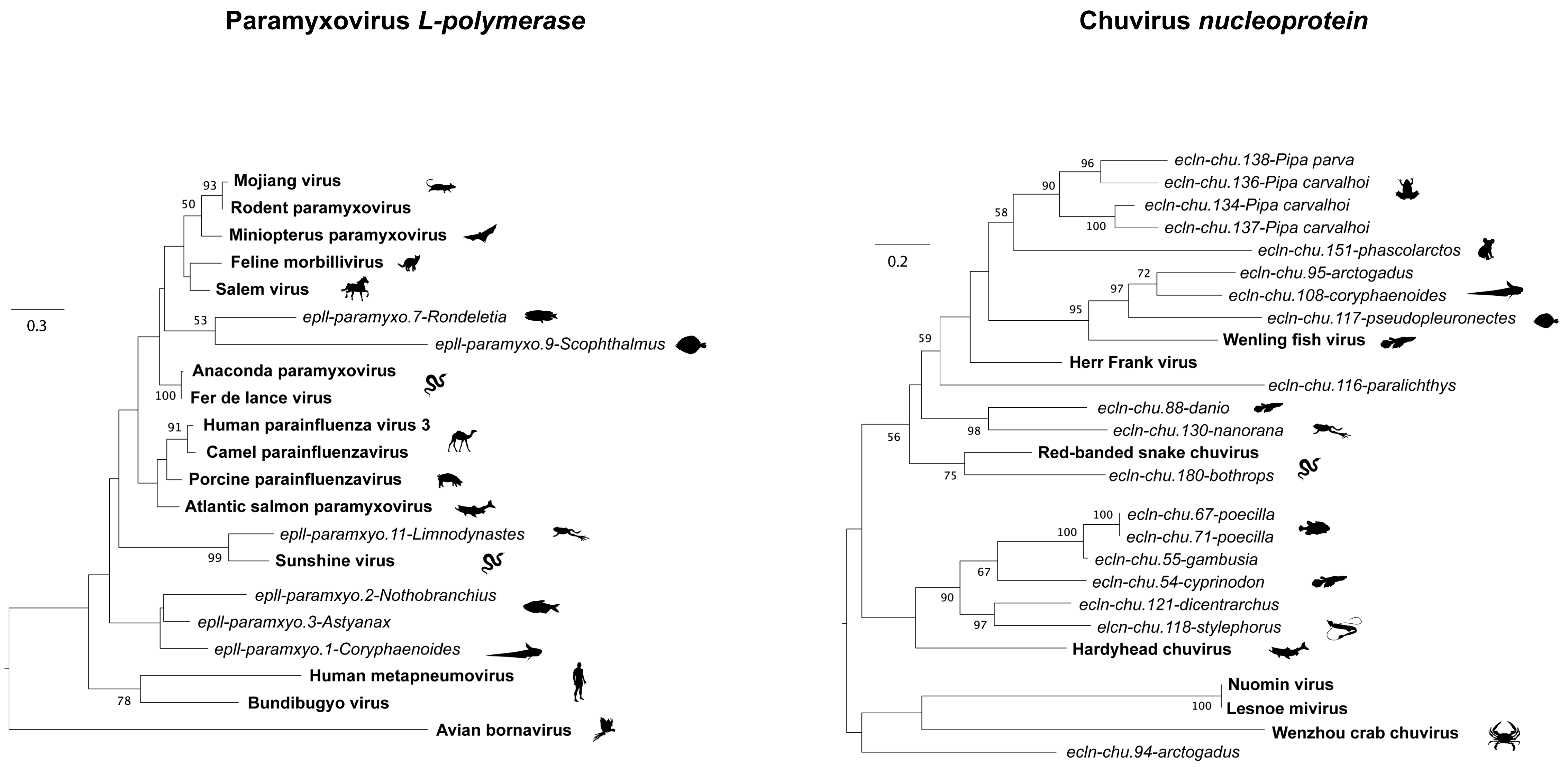
Paramyxovirus L-polymerase
Mojiang virus
Rodent paramyxovirus
Miniopterus paramyxovirus
Feline morbillivirus
Salem virus
epII-paramyxo.7-Rondeletia
epII-paramyxo.9-Scophthalmus
Anaconda paramyxovirus
Fer de lance virus
Human parainfluenza virus 3
Camel parainfluenzavirus
Porcine parainfluenzavirus
Atlantic salmon paramyxovirus
epII-paramxyo.11-Limnodynastes
Sunshine virus
epII-paramxyo.2-Nothobranchius
epII-paramxyo.3-Astyanax
epII-paramxyo.1-Coryphaenoides
Human metapneumovirus
Bundibugyo virus
Avian bornavirus
0.3
93
50
53
100
91
99
78
Chuvirus nucleoprotein
ecln-chu.138-Pipa parva
ecln-chu.136-Pipa carvalhoi
ecln-chu.134-Pipa carvalhoi
ecln-chu.137-Pipa carvalhoi
ecln-chu.151-phascolarctos
ecln-chu.95-arctogadus
ecln-chu.108-coryphaenoides
ecln-chu.117-pseudopleuronectes
Wenling fish virus
Herr Frank virus
ecln-chu.116-paralichthys
ecln-chu.88-danio
ecln-chu.130-nanorana
Red-banded snake chuvirus
ecln-chu.180-bothrops
ecln-chu.67-poecilla
ecln-chu.71-poecilla
ecln-chu.55-gambusia
ecln-chu.54-cyprinodon
ecln-chu.121-dicentrarchus
elcn-chu.118-stylephorus
Hardyhead chuvirus
Nuomin virus
Lesnoe mivirus
Wenzhou crab chuvirus
ecln-chu.94-arctogadus
0.2
96
90
100
58
72
97
95
59
98
56
75
100
100
67
90
97
100

**Figure S7g-h.**

#### Bornavirus *L-polymerase*

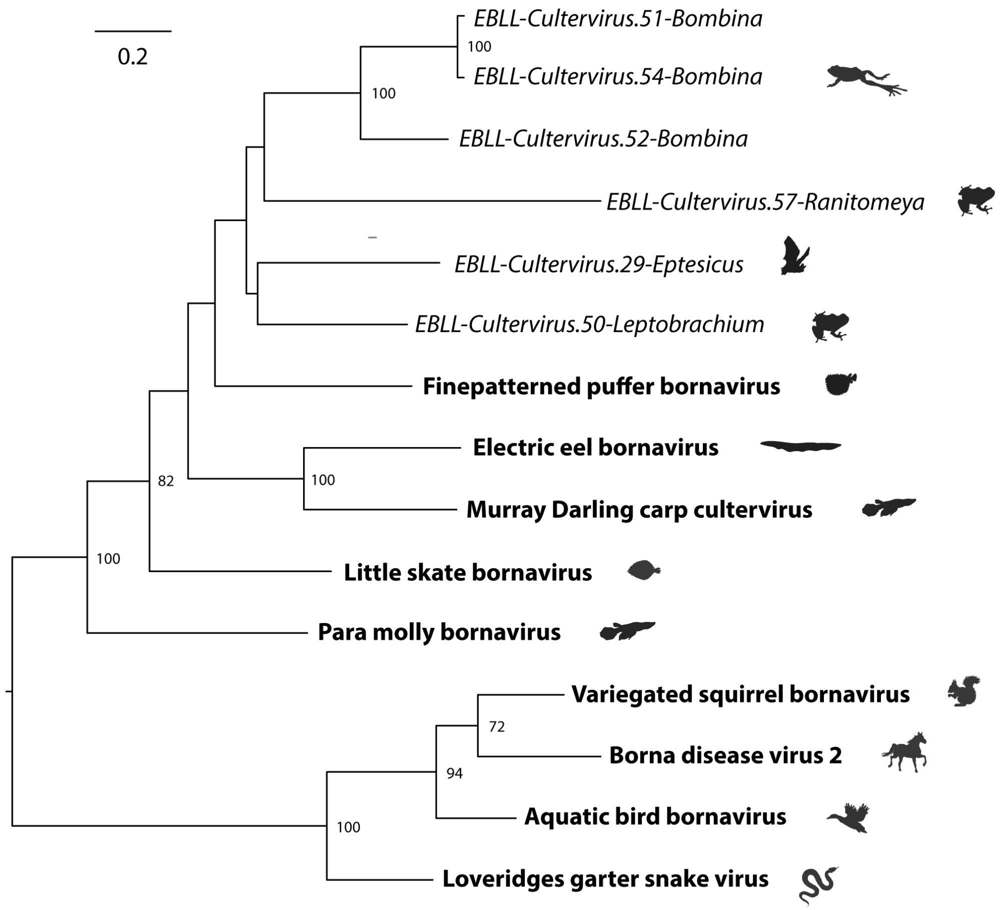


#### Bornavirus *glycoprotein*

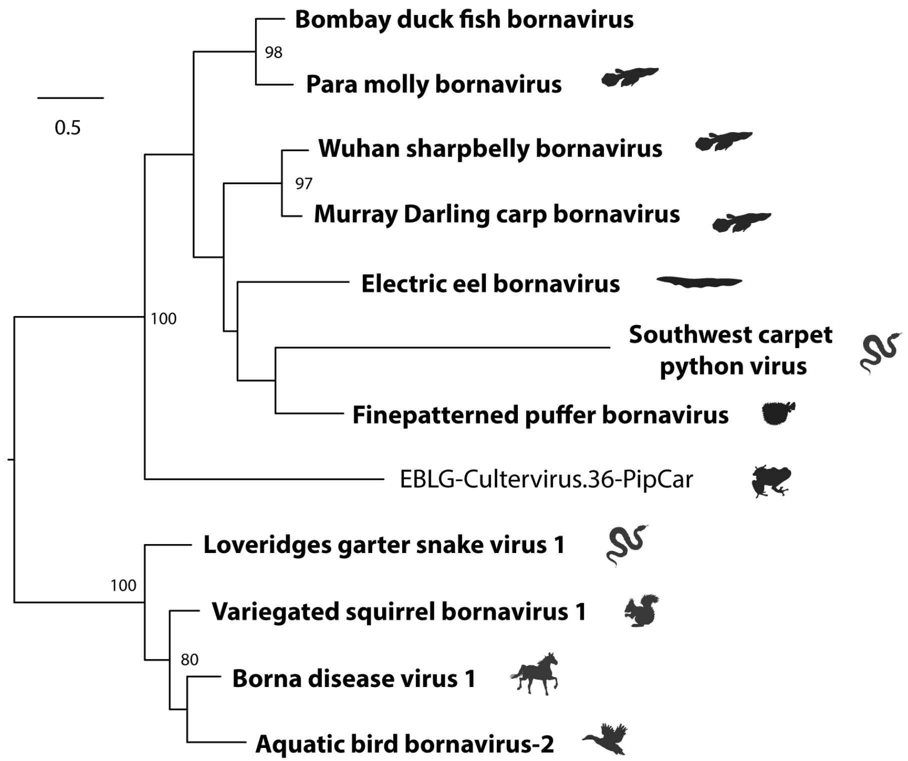


**Figure S7i-j.**

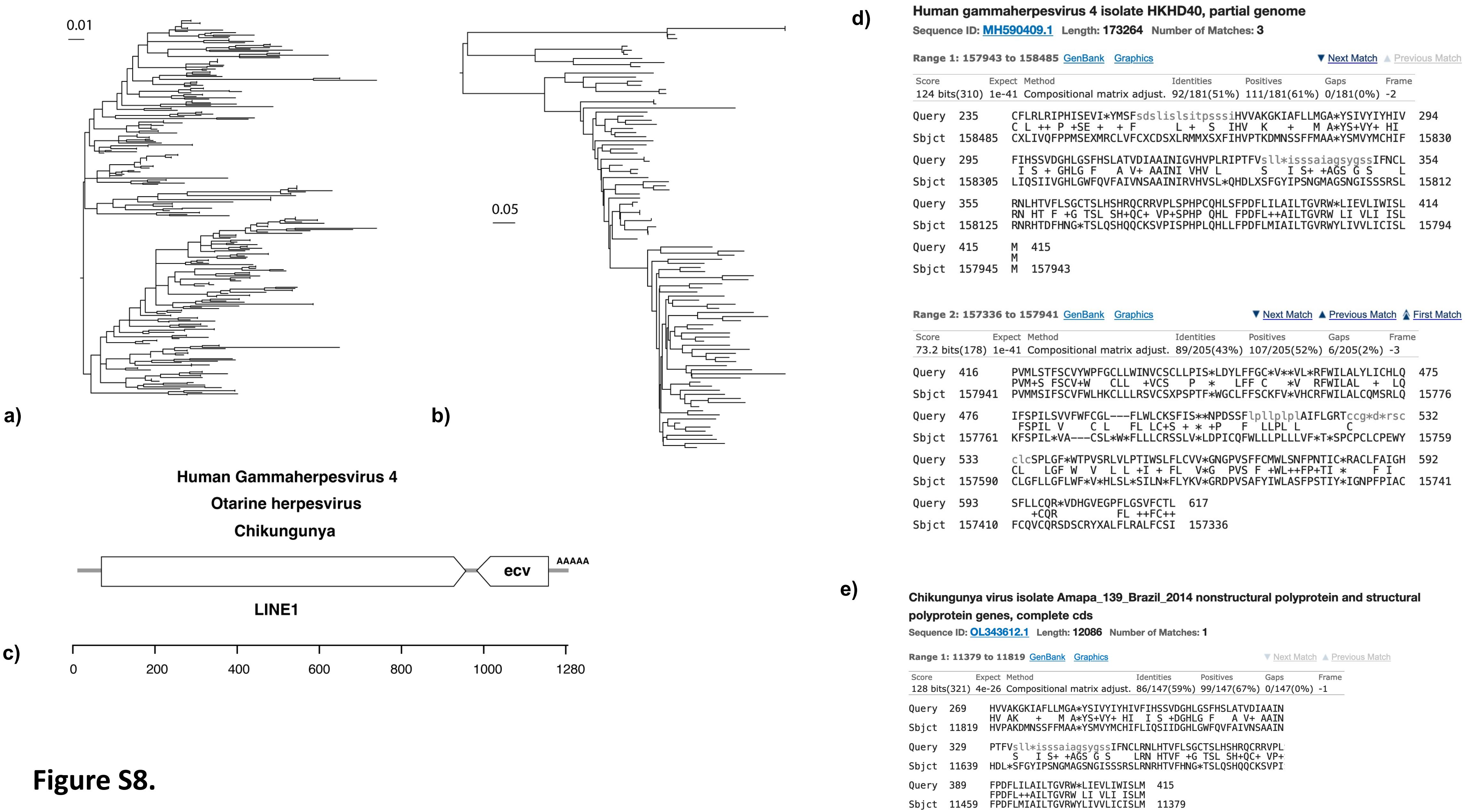
0.01
a)
0.05
b)
Human Gammaherpesvirus 4
Otarine herpesvirus
Chikungunya
ecv
AAAAA
LINE1
c)
0
200
400
600
800
1000
1280
d)
Human gammaherpesvirus 4 isolate HKHD40, partial genome
Sequence ID: MH590409.1 Length: 173264 Number of Matches: 3
Range 1: 157943 to 158485 GenBank Graphics
Next Match Previous Match
Score Expect Method Identities Positives Gaps Frame
124 bits(310) 1e-41 Compositional matrix adjust. 92/181(51%) 111/181(61%) 0/181(0%) -2
Query 235 CFLRLRIPHISEVI*YMSFsdslislsitpsssiHVVAKGKIAFLLMGA*YSIVYIYHIV 294
C L ++ P +SE + + F L + S IHV K + M A*YS+VY+ HI
Sbjct 158485 CXLIVQFPPMSEXMRCLVFCXCDSXLRMMXSXFIHVPTKDMNSSFFMAA*YSMVYMCHIF 15830
Query 295 FIHSSVDGHLGSFHSLATVDIAAINIGVHVPLRIPTFVsll*isssaiagsygssIFNCL 354
I S + GHLG F A V+ AAINI VHV L S I S+ +AGS G S L
Sbjct 158305 LIQSIIVGHLGWFQVFAIVNSAAINIRVHVSL*QHDLXSFGYIPSNGMAGSNGISSSRSL 15812
Query 355 RNLHTVFLSGCTSLHSHRQCRRVPLSPHPCQHLSFPDFLILAILTGVRW*LIEVLIWISL 414
RN HT F +G TSL SH+QC+ VP+SPHP QHL FPDFL++AILTGVRW LI VLI ISL
Sbjct 158125 RNRHTDFHNG*TSLQSHQQCKSVPISPHPLQHLLFPDFLMIAILTGVRWYLIVVLICISL 15794
Query 415 M 415
M
Sbjct 157945 M 157943
Range 2: 157336 to 157941 GenBank Graphics
Next Match Previous Match First Match
Score Expect Method Identities Positives Gaps Frame
73.2 bits(178) 1e-41 Compositional matrix adjust. 89/205(43%) 107/205(52%) 6/205(2%) -3
Query 416 PVMLSTFSCVYWPFFGCLLWINVCSCLLPIS*LDYLFFGC*V**VL*RFWILALYLICHLQ 475
PVM+S FSCV+W CLL +VCS P * LFF C *V RFWILAL + LQ
Sbjct 157941 PVMMSIFSCVFWLHKCLLLRSVCSXPSPTF*WGCLFFSCKFV*VHCRFWILALCQMSRLQ 15776
Query 476 IFSPILSVVFWFCGL---FLWLCKSFIS**NPDSSFlpllplplAIFLGRTccg*d*rsc 532
FSPIL V C L FL LC+S + * +P F LLPL L C
Sbjct 157761 KFSPIL*VA---CSL*W*FLLLCRSSLV*LDPICQFWLLLPLLLVF*T*SPCPCLCPEWY 15759
Query 533 clcSPLGF*WTPVSRLVLPTIWSLFLCVV*GNGPVSFFCMWLSNFPNTIC*RACLFAIGH 592
CL LGF W V L L +I + FL V*G PVS F +WL++FP+TI * F I
Sbjct 157590 CLGFLLGFLWF*V*HLSL*SILN*FLYKV*GRDPVSAFYIWLASFPSTIY*IGNPFPIAC 15741
Query 593 SFLLCQR*VDHGVEGPFLGSVFCTL 617
+CQR FL ++FC++
Sbjct 157410 FCQVCQRSDSCRYXALFLRALFCSI 157336
e)
Chikungunya virus isolate Amapa_139_Brazil_2014 nonstructural polyprotein and structural polyprotein genes, complete cds
Sequence ID: OL343612.1 Length: 12086 Number of Matches: 1
Range 1: 11379 to 11819 GenBank Graphics
Next Match Previous Match
Score Expect Method Identities Positives Gaps Frame
128 bits(321) 4e-26 Compositional matrix adjust. 86/147(59%) 99/147(67%) 0/147(0%) -1
Query 269 HVVAKGKIAFLLMGA*YSIVYIYHIVFIHSSVDGHLGSFHSLATVDIAAIN
HV AK + M A*YS+VY+ HI I S +DGHLG F A V+ AAIN
Sbjct 11819 HVPAKDMNSSFFMAA*YSMVYMCHIFLIQSIIDGHLGWFQVFAIVNSAAIN
Query 329 PTFVsll*isssaiagsygssIFNCLRNLHTVFLSGCTSLHSHRQCRRVPL
S I S+ +AGS G S LRN HTVF +G TSL SH+QC+ VP+
Sbjct 11639 HDL*SFGYIPSNGMAGSNGISSSRSLRNRHTVFHNG*TSLQSHQQCKSVPI
Query 389 FPDFLILAILTGVRW*LIEVLIWISLM 415
FPDFL++AILTGVRW LI VLI ISLM
Sbjct 11459 FPDFLMIAILTGVRWYLIVVLICISLM 11379

Figure S8.
